# Trait evolution with incomplete lineage sorting and gene flow: the Gaussian Coalescent model

**DOI:** 10.64898/2026.03.10.710880

**Authors:** Cécile Ané, Paul Bastide

## Abstract

Most phylogenetic comparative methods use a species-level phylogeny, ignoring the effect of incomplete lineage sorting (ILS) and hemiplasy on the traits of interest. We consider here a trait controlled additively by one or more unknown loci. Their gene trees may differ from the species phylogeny due to ILS. The species phylogeny can be any phylogenetic tree or network or admixture graph on any number of taxa or populations. Our process of trait evolution accounts for introgression represented in the phylogeny by hybrid (or admixture) edges, which capture gene flow, admixture or hybridization events. The process models the trait in each individual, for any number of individuals from each taxon. Our model allows for polymorphism in the ancestral population at the root of the species phylogeny, and we can derive the heritable within-population variation expected from our model. Even if each locus evolves according to a Brownian motion, the joint distribution of the trait across all measured individuals is not generally Gaussian due to ILS. We provide a Gaussian approximation, named the Gaussian Coalescent (GC), and show how to compute its variance matrix efficiently using a single traversal of the species phylogeny. In simulations, this model is much more accurate than the model ignoring ILS. In simulations and on a data set of tomato floral traits, it is favored over the standard Brownian motion model with extra within-population variance. The GC model opens new avenues for various phylogenetic methods, accounting for hemiplasy and gene flow simultaneously on any general network or admixture graph. It is implemented in `phylolm` v2.7.0 and in `PhyloTraits` v1.2.0.

## 1. Introduction

Phylogenetic comparative methods (PCMs) leverage the historical relationships between taxa of interest to discover patterns of trait evolution. They have a vast array of applications in evolutionary biology (Felsenstein, 2004; Harmon, 2019; Revell and Harmon, 2022). Traditionally, the evolution of a trait is modeled along the species phylogeny. Only recently was it recognized that spurious conclusions may arise in the presence of gene-tree discordance, when individual genes have genealogies that differ from the species phylogeny. For binary traits, hemiplasy (Avise and Robinson, 2008) is a source of trait incongruence relative to the species phylogeny, when a single transition on a discordant gene tree can explain the trait pattern whereas homoplasy would instead require several convergent transitions along the species phylogeny, either a tree (Guerrero and Hahn, 2018) or a network with reticulations (Hibbins et al., 2020; Wang et al., 2021). Incomplete lineage sorting (ILS) is one of the main identified biological causes for gene-tree discordance (Degnan and Rosenberg, 2009; Maddison, 1997; Schrempf and Szöllösi, 2020). Within a population, it can be modeled by the coalescent process (Kingman, 1982). When applied to each population of a phylogenetic tree or network, this process is typically called the multispecies coalescent (MSC, Degnan, 2018; Rannala and Yang, 2003). On a network, parental inheritance can be assumed independent, correlated or even shared across lineages in admixed populations (Fogg et al., 2023).

Continuous polygenic traits are controlled by a large number of genes, each of which being possibly affected by ILS. Classical models use a single stochastic process such as the Brownian motion (BM) on the species tree (Felsenstein, 1973) or network (Bastide et al., 2018). These models induce phylogenetic correlation, which can then be accounted for in statistical analyses such as linear regression or ANOVA, to avoid the detection of spurious effects if ignored (Felsenstein, 1985). However, traditional models do not account for the diverse histories of the gene trees underlying the trait. Mendes et al. (2018) pioneered the use of the MSC in this context. They assume that the measured trait *X* is the sum of the effects *Y* ^(*l*)^ of *L* genes, each following its own stochastic process on its own gene tree. Under the MSC, they derived theoretical formulas for the covariance matrix of such a trait measured at the tips of a phylogenic tree, and provided a closed-form expression for triplets. Hibbins et al. (2023) built on this approach and implemented the computation of a triplet-based approximation for trees with branches scaled in coalescent units in the R package `seastaR`, and demonstrated its use on a wild tomato flower trait dataset. Their work underlined the importance of taking ILS into account in PCMs. Their method has some limitations: it requires one trait value per population, does not consider within-population variation, and cannot be used on a phylogenetic network. More importantly, it is based on conditioning the evolution of each effect *Y* ^(*l*)^ to its value at the root of its own gene tree, which can be arbitrarily far in the past. As we show below, this induces some unexpected behavior of the model, that, for a phylogeny with a given root population, can predict a different covariance between the trait of two populations *A* and *B* whether a third population *C* is included in the study or not.

In this work, we propose a general model for the evolution of a polygenic trait on a species network with ILS. We assume that the gene trees are distributed in the phylogeny according to the network multispecies coalescent process with independent inheritance of parental lineages at reticulations (Degnan, 2018; Fogg et al., 2023). As in Mendes et al. (2018), we assume that the continuous trait is the sum of the effects of many genes, each evolving on its own gene tree, but we use a different conditioning at a fixed point in time, at or before the root of the phylogeny, to ensure a consistent model across genes. We derive the covariance matrix of the univariate trait, and show that the distribution of the vector of measured traits converges under mild conditions to a multivariate Gaussian. We coin this distribution the “Gaussian-Coalescent” (GC). The computation of the GC covariance matrix involves one traversal of the phylogeny, and has an explicit closed form in the tree case. It depends on two parameters: the rate of evolution per coalescent unit 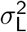, and the trait variance in the ancestral population v_0_. While classical BM models only describe the evolution of the population mean (Felsenstein, 2004), we derive formulas for how both the mean and the variance of the trait within each population propagate along the phylogeny. This implies that the GC can directly give an expectation of the within-population variance on the phylogeny, that can be compared to the classical BM model with additional within-population variation (Teo et al., 2023). Further, our model recovers several classical frameworks: the standard BM when ILS disappears, a single population when ILS dominates, and the evolution of an allele frequency when considering a single locus and no mutation.

In the remainder, we most often refer to taxa as “populations”, but our model and theory equally applies to species-level phylogenies. We keep referring to “species tree” and “species networks” because these terms are frequently used in studies of the coalescent. We also refer to the genealogy of any locus as a “gene tree”, even for a single non-coding position in the genome, also to follow conventional terminology. The paper is organized as follows. In Section 2, we formally define our model of polygenic trait evolution under ILS (Sec. 2.1), give exact recursive formulas for computing its moments on a phylogenetic network (Sec. 2.2), show how the model can be re-written as a sum of independent Brownian motions on re-scaled phylogenies (Sec. 2.3), and explore its links to existing models in the admixture graph literature (Sec. 2.4), before introducing the Gaussian-Coalescent model (Sec. 2.5) and showing how it differs from previous PCM approaches (Sec. 2.6). Most of the mathematical proofs are in the Supplementary Material (SM). We then study the GC model in Section 3, detailing its implementation (Sec. 3.1), its behavior on simulated data (Sec. 3.2) and on wild tomato floral traits (Sec. 3.3). Section 4 concludes with a discussion and some recommendations.

## 2. Continuous trait evolution model with ILS

In this section, we define a process-based model for the evolution of a polygenic trait on a phylogeny (Sec. 2.1), and then show how to compute the expectation vector and the variance matrix from this process on a phylogeny (Sec. 2.2). The core Gaussian-Coalescent (GC) model is summarized in Section 2.5. Sections 2.3, 2.4 and 2.6 (marked with ^∗^) describe some of the properties of the model and its links with previous approaches from population genetics and phylogenetic comparative methods, and can be skipped to focus on the definition and use of the model.

Throughout, we use the standard definition of the variance matrix: the variance of the trait measured on one individual is defined as the expected squared deviation from the expected value of the trait under the model, and the covariance between the trait of two individuals is the expected value of the product of their deviations from their respective expected value. The phylogenetic variance matrix (also called the variance - covariance matrix) contains the trait co-variance for each pair of measured individuals.

### 2.1. Polygenic evolution model on a phylogeny

We start by describing a model for the evolution of a polygenic trait *X* within a single population, and then show how to extend the model to a phylogeny. We refer to Felsenstein (2004, Chap. 24) for background on quantitative characters.

#### 2.1.1. Polygenic evolution model in one population

We assume that the trait is determined by *L* genetic loci, each having an infinitesimally small effect *Y* ^(*l*)^ and evolving according to the following process. For simplicity, we also assume haploid individuals, each carrying one copy of each locus.

##### Definition 1

(Lévy evolution of the effect of one locus *l*). Let *l* be a locus, with a gene genealogy *G*_*l*_ . We assume that *G*_*l*_ is drawn from the coalescent process (backwards in time) and, as in Mendes et al. (2018), that the evolution of the locus effect *Y* ^(*l*)^ along *G*_*l*_ (forward in time) is independent of the realization of the coalescent process that generated *G*_*l*_, as under neutral evolution in the absence of selection. The effect *Y* ^(*l*)^ is assumed to evolve according to a Lévy process *D*_*θ*_ with parameter *θ* along *G*_*l*_, as in Landis et al. (2013). Lévy processes have independent stationary increments: for any 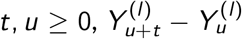 is independent of 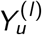 and has distribution *D*_*θ*_(*t*). Then, within each edge in *G*_*l*_, the change in 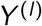 over *t* units of time has distribution *D*_*θ*_(*t*), and the process splits into two independent processes starting at the same value at each node of the tree *G*_*l*_ . We also assume that the process has mean 0 (centered) and finite variance 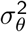 at time 1, so that *D*_*θ*_(*t*) is also centered for any *t*, and with variance 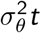.

Below are two common examples of processes that satisfy Def. 1.

##### Example 1

(Brownian motion). The Brownian motion with variance rate *σ*^2^ is the most common example of a centered Lévy process, with 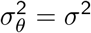. A Brownian motion with a trend is also a Lévy process, but not centered.

##### Example 2

Mutational model). A traditional mutational model assumes that changes occur in *Y* ^(*l*)^ via mutations, with mutation rate *µ* per locus per generation, and individual mutational effects (jumps in *Y* ^(*l*)^) drawn from a distribution with mean 0 and variance 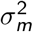 (used in e.g., Schraiber and Landis, 2015). Then *Y* ^(*l*)^ evolves according to a compound Poisson process, which is a centered Lévy process with variance rate 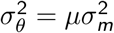.

Other examples include processes studied by Landis et al. (2013) like the variance-gamma process and the symmetric *α*-stable processes (including the Cauchy process), although some *α*-stable processes have infinite variance (at any time *t*), so that they do not enter in our present setting (Schraiber and Landis, 2015). Assuming additive effects, this locus effect model leads to the following model for the trait *X* .

##### Definition 2

(Polygenic model in one population). We assume that each locus *l* follows the model in Def. 1, and has an effect *Y* ^(*l*)^ on the trait, acting additively across loci:

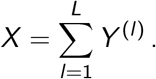

We also assume loci are independent. This corresponds to free recombination between loci (for independent gene trees) and selective neutrality (for independent effects given the gene trees, e.g. Bulmer, 1971). Then the phenotype *X* is itself a Lévy process as a sum of independent Lévy processes, with a finite rate

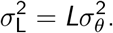

#### 2.1.2. Phylogenetic model

We now turn to the case when several populations are observed. These populations are related by a given phylogeny *N*, which may have reticulations, and we define a model for the evolution of the trait on this phylogeny.

##### Definition 3

(Phylogenetic model for one locus). Let *N* be a known phylogenetic network specifying the relationships between the populations from which individuals are sampled, such that each of its edges *e* has length *ℓ*_*e*_ in *coalescent units*. Let *l* be a given locus, with a gene genealogy *G*_*l*_ drawn from the network multispecies coalescent process. As in Def. 1, we assume that the evolution of the locus effect *Y* ^(*l*)^ along *G*_*l*_ is independent of the realization of the coalescent process that generated *G*_*l*_ . Further, we assume that there is a unique ancestral population. A standard choice is the most recent common ancestor (MRCA) of all sampled populations when *N* is a tree, or the lowest stable ancestor more generally (Steel, 2016). Without loss of generality, we may fix this ancestral population to be the root *ρ* of *N*, possibly attached to a single “root edge”. The evolution of the locus effect on *N* is then as follows.

- At the root population *ρ*, the effect *Y* ^(*l*)^ of ancestral individuals in *G*_*l*_ are drawn independently from a common distribution *P*_0_, with mean *m*_0_*/L* and variance v_0_*/L*.
- Along the gene tree *G*_*l*_, *Y* ^(*l*)^ evolves according to a centered Lévy process with finite variance as in Def. 1.
- The parameter 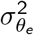 of the process may differ across branches of *N* .

*Remark* 1 (Population size and time in generations). Since the phylogeny *N* has branch lengths in coalescent units, 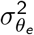 is a variance rate per coalescent unit so that 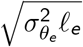 has the same unit as the trait being studied (for instance, if the trait is a corolla diameter in millimeters, then 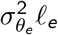 would be in mm^2^). If one has information on the haploid effective population size 2*N*(*e*) in branch *e*, then the branch length in generations can be recovered as *κ*_*e*_ = 2*N*(*e*)*ℓ*_*e*_, and the rate of evolution per generation is given by 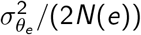. In the following, we make no assumption on effective population sizes, which can vary over time. The rate 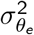 can vary across branches, to account for an accelerating or decelerating rate of evolution per coalescent unit. If population sizes are known, different assumptions on the rate of evolution in number of generations can be used. Combining Defs. 2 and 3, we get an evolutionary model for the measured polygenic trait *X* .

##### Definition 4

(Phylogenetic polygenic model). A trait *X* follows a phylogenetic polygenic model on a phylogenetic network *N* if it is the sum of independent additive effects *Y* ^(*l*)^ from *L* loci, each following the model of Def. 2 on *N* . In particular, the distribution *P*_0_ of *X* in the root population *ρ* has mean *m*_0_ and variance v_0_, and the rate of evolution on each branch *e* of *N* is given by 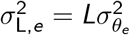.

##### Definition 5

(Equilibrium variance ratio). We define the equilibrium variance ratio as

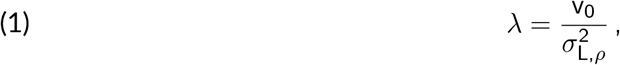

where 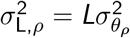 is the equilibrium trait variance, from an ancestral lineage that evolved according to a Lévy process with variance 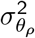 at each locus (Lynch and Hill, 1986). The root population *ρ* is at equilibrium when *λ* = 1.

To derive the expectation and variance matrix of the trait values from the process in Def. 4, we will need coalescence probabilities and expected times, as recalled below (from Mendes and Hahn 2018, Eq. (A.7) in Supplementary Material on dryad Mendes and Hahn, 2017).

##### Lemma 1

(Coalescent probability and shared time). *For a time ℓ in coalescent units, define:*

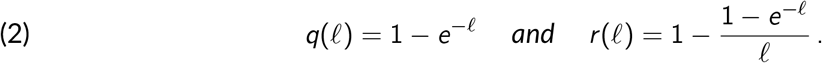

*Two individuals from the same population coalesce within time ℓ with probability q*(*ℓ*). *In other words q*(*ℓ*) = ℙ [*T < ℓ*] *where T is their random coalescent time. These two individuals share a common ancestor for an average of ℓr* (*ℓ*) *coalescent units within the past ℓ coalescent units, that is:* E [(*ℓ* − *T*)_+_] = *ℓr* (*ℓ*). *Both q*(*ℓ*) *and r* (*ℓ*) *are between* 0 *and* 1, *are* 0 *at ℓ* = 0, *and tend to* 1 *when ℓ* = ∞.

### 2.2. Covariances on a species network for a single locus effect

In this section, we derive explicit formulas to compute the covariance between the traits measured at various nodes and individuals in the phylogeny, assuming the model of Def. 3 to focus on a single locus effect *Y* . From Def. 4 and by additivity of variances for independent variables, the moments for the polygenic trait *X* simply follow. Compared to Mendes et al. (2018) and subsequent papers based on their calculations (e.g., Hibbins et al., 2023), we clarify the conditioning and parameters when taking expectations, variances and covariances, and we explicitly model ancestral polymorphism.

In this section, we state our results and provide informal justifications. The formal proofs rely on the definition of a graphical model, and are in SM Section A.

#### 2.2.1. Covariances on a network for a single locus effect

We assume that *Y* evolves according to the model in Def. 3 on a species phylogeny *N*, which may have reticulations. To specify the full variance-covariance matrix of *Y*, we use the following quantities.

##### Definition 6

(Expectation, variance and covariances). Take any two nodes *u* and *v* in the phylogeny *N*. We write *E*_*u*_ the expectation and Φ_*u*_ the total variance of the trait *Y* of a random single individual in the given population *u*. The covariance between two *different* random individuals taken from populations *u* and *v* is denoted by Ω_*u,v*_ (when *u* = *v*, we take two different individuals from the same population *u*). To simplify notations, we omit *ρ*, but all expectations, variances and covariances are calculated conditional on the trait distribution in *ρ*.

These notations and their link to the coalescent are illustrated in Fig. 1. The following result shows that **E, Φ** and **Ω** can be computed recursively with one preorder traversal of the phylogeny.

**Figure 1.**
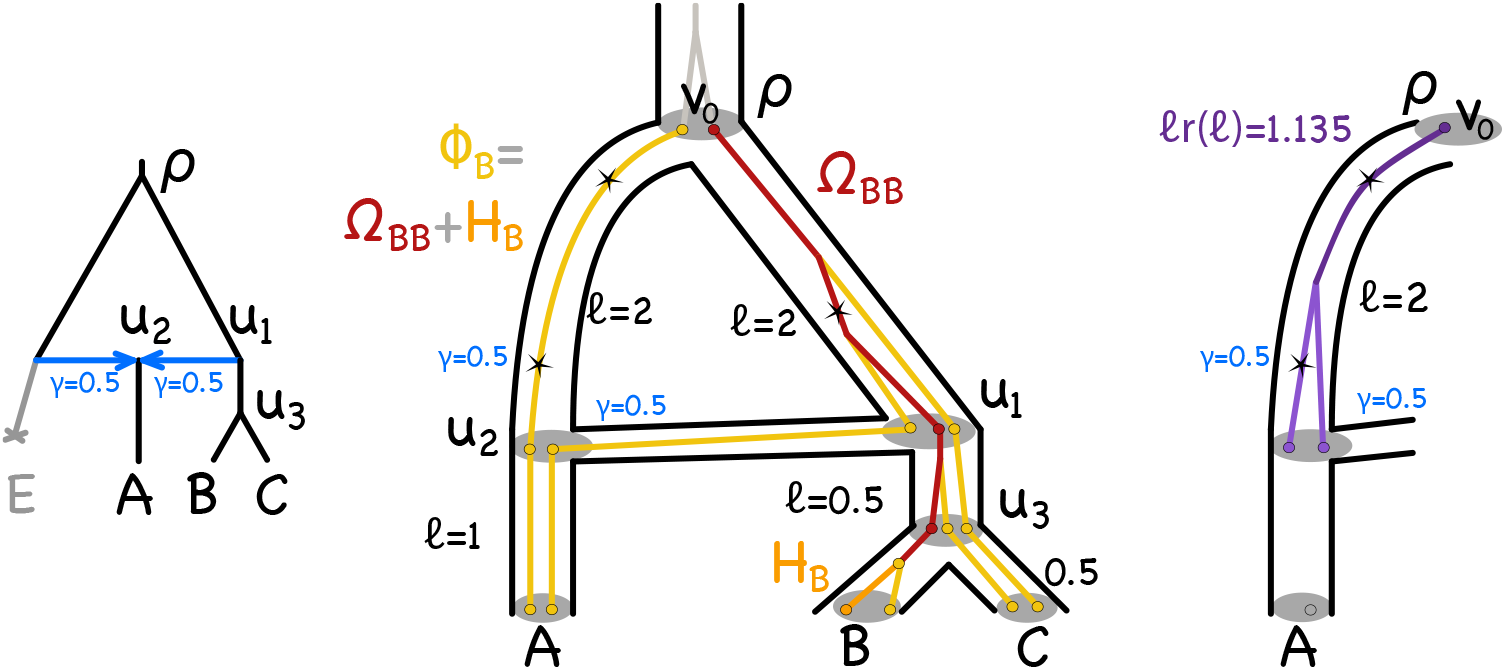
Network to illustrate the variances Φ_*u*_ (for 1 random individual from *u*) and covariances Ω_*u,v*_ (for 2 random individuals from *u* and *v*) conditional on the ancestral population *ρ*. Expectations and variances are taken over all the possible gene trees. **Left**: population history with *u*_2_ arising from hybrid speciation with proportion *γ* = 0.5 of its genome from each parent, and *E* either extinct or unsampled. **Middle**: phylogenetic network (black) for *A, B* and *C* with 2 individuals sampled from each, and an example gene tree for one sampled locus (colored). The locus effect evolves on the gene tree according to a centered Lévy process (Def. 1), such as a compound Poisson process with mutations (black stars) appearing at rate *µ* and adding an effect of variance 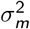 on the trait, with total variance rate 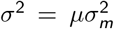. Ancestral populations, in preorder, are *ρ, u*_1_, *u*_2_ and *u*_3_. Ω_*B,B*_ is the expected time of shared ancestry for 2 individuals randomly sampled from *B* (in red in the example gene tree) up to *ρ*, scaled by the evolutionary rate *σ*^2^, plus a term that depends on the root variance v_0_. Φ_*B*_ is the scaled expected time from a random individual in *B* up to *ρ*, plus v_0_. The difference *H*_*B*_ = Φ_*B*_ − Ω_*B,B*_ is the expected within-population variance within *B* (Lemma 7), and is the scaled expected time between the individual in *B* and the first coalescent event (orange line), plus a term that depends on v_0_. In the example gene tree, the 2 individuals from *A* do not coalesce along the lineage from *u*_2_ to *A*, and their ancestors in *u*_2_ are inherited from different parental populations. **Right**: alternative scenario, in which both individuals in *u*_2_ are inherited from the same parental population. The formulas involve *ℓr* (*ℓ*), the expected time of shared ancestry within a lineage of *ℓ* coalescent units (dark purple in the example gene tree).

##### Proposition 2

(Recursion formula). *Let N be a species phylogeny. For an edge e in N, let ℓ*_*e*_ *denote its length in coalescent units. Assume that Y evolves according to the model of Def. 3 on N, with increments of finite variance rate* 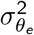. *The mean and variance of Y among individuals in ρ, the root population, are denoted by m*_0_ *and* v_0_. *Then for any node u of N*,

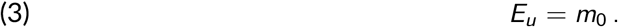

*At the root*, Φ_*ρ*_ = v_0_ *and* Ω_*ρ,ρ*_ = 0. *Let u and v be two nodes in the network, such that v is* not *a descendant of u. Such is the case if v is listed before u in preorder. If u is a tree node with parent p and parent edge e then*

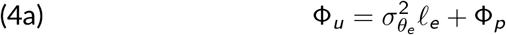

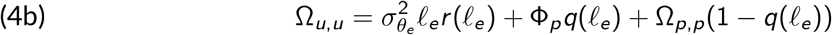

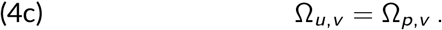

*More generally, if u has m parents p*_1_, …, *p*_*m*_, *with edge e*_*k*_ = (*p*_*k*_, *u*) *of variance rate* 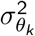, *length ℓ*_*k*_ *in coalescent units and inheritance γ*_*k*_, *then*

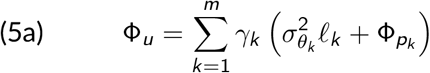

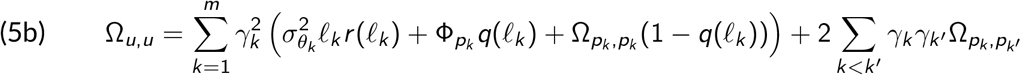

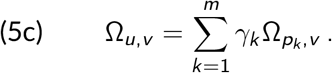

*Proof*. The formal proof is in SM Section A. Informally, in the tree case, for a single individual in a single population, (3) and (4a) directly follow from the assumption of centered Lévy increments with finite variance 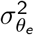, while (4c) comes from the conditional independence structure given by the phylogeny along with the preorder of the nodes. In (4b), the three terms can be interpreted using Lemma 1. The first term is just the variance of the process 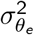 times *ℓ*_*e*_*r* (*ℓ*_*e*_), which is the expected time that the two different individuals from population *u* share a common ancestor within the branch of length *ℓ*_*e*_. Forward in time, the two individuals shared the same trait for this amount of time, and then split, following independent processes. If these two individuals do coalesce within the branch (backwards in time), then they inherit from a single individual in the parent population *p* with variance Φ_*p*_. This happens with probability *q*(*ℓ*_*e*_), giving rise to the second term. Finally, with probability 1 − *q*(*ℓ*_*e*_), the two individuals do not coalesce within the branch, and inherit from the covariance Ω_*p,p*_ of two distinct individuals within parent population *p*, which gives the third term. General formulas (5) have similar interpretations, but allow for node *u* to have several parent populations. □

The next proposition states that the recursion formulas of Prop. 2 are robust to sampling: the covariance between two individuals sampled from two populations does not depend on whether a third individual is sampled or not. On the network in Fig. 1 for example, Ω_*A,B*_ is the same whether *E* is sampled or not. This property, which directly stems from our process-based definition of the model, makes it different from the formulas in Mendes et al. (2018), that are not robust to sub-sampling in general (see Section 2.6).

##### Proposition 3

(Sampling stability). *Assume a model on a network N such that the covariances are given by Prop. 2 above. For a taxon t, define N*_−*t*_ *the pruned subnetwork of N with t removed, but keeping the same root (possibly leaving the root connected by a single branch to the most recent stable ancestor of the remaining leaves) and not suppressing degree-2 nodes. Then, for any population nodes u and v that remain in N*_−*t*_ *(possibly with u* = *v)*, Φ_*u*_ *and* Ω_*u,v*_ *given by Prop. 2 are the same when the recursion is applied to N*_−*t*_ *as when it is applied to the full network N. Further, if the rate* 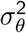 *is constant across branches, then the non-root degree-2 nodes can be suppressed from N*_−*t*_ *without altering the distribution of the remaining nodes*.

The proof of this result is presented SM Section A. Note that it is generally useful to allow for degree-2 nodes in the phylogeny to allow for changes in evolutionary parameters, such as bottlenecks or other changes in population sizes, or changes in the rate of phenotypic evolution. These parameters *N*(*e*) and 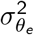 can vary across edges but are fixed within an edge *e*. To model variation in *N* or 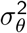 within an edge *e* from *u* → *v*, one may subdivide *e* into edges *e*_*1*_ from *u* → *w* and *e*_*2*_ from *w*→*v*, where *w* is a new node of degree-2. This brings more flexibility in the model, by allowing for *N*(*e*_1_)≠ *N*(*e*_2_) or 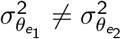.

**Example: using the recursive (co)variance calculations on the network in Fig. 1**

We illustrate Prop. 2 assuming a constant evolutionary rate 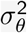 across branches.

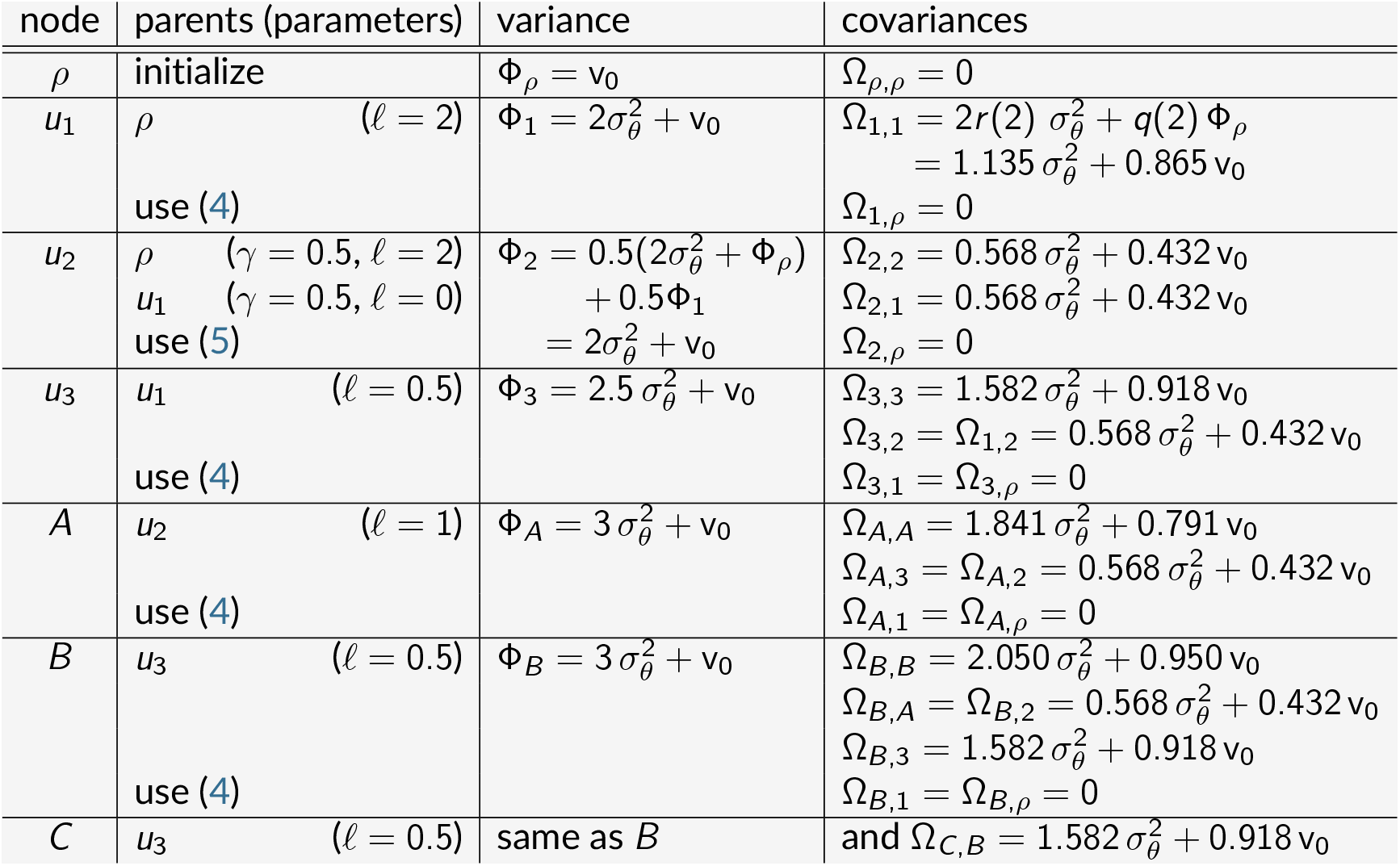

Regardless of 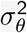 and v_0_, we get that Ω_*A,A*_ *<* Ω_*B,B*_. This is because *A* is below a reticulation: 2 individuals in *A* have reduced covariance as one might inherit from gene flow and the other not. From Lemma 7, this implies that the within-population variance is expected to be greater in *A* than in *B*. Here we obtain Φ_*A*_ = Φ_*B*_ = Φ_*C*_ because the network is time-consistent (all paths from *ρ* to *A* have the same length) and ultrametric (all paths from *ρ* to *A, B* or *C* have the same length).

### 2.3. Additivity and branch length transformation^∗^

In this section, we explore two interesting properties of the model, and show that it can be seen as a sum of two independent Brownian motions on phylogenies with modified branch lengths. This section can be skipped to focus on the definition and use of the model.

#### 2.3.1. Additivity of mutation and drift

We can decompose the effect of one locus *Y* as the sum of two independent processes: one representing trait evolution through molecular change after *ρ* (modeled by a Lévy process), and one representing pure drift through the coalescent. For example, all terms calculated in the example on network in Fig. 1 are linear combinations of 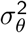 (the Lévy contribution) and of v_0_ (from drift and ancestral polymorphism in *ρ*). Indeed, the recursion formulas of Prop. 2 can be re-written as follows.

##### Corollary 4

(Additivity). *Using the same model and notations as in Prop. 2, we can decompose*

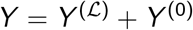

*with Y* ^(ℒ)^ *and Y* ^(0)^ *independent, Y* ^(ℒ)^ *the process following Def. 3 with m*_0_ = 0 *and* v_0_ = 0, *and Y* ^(0)^ *the process following Def. 3 with the same m*_0_ *and* v_0_ *as Y, but* 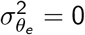 *on all branches. In other words, Y* ^(ℒ)^ *captures the evolutionary Lévy process after ρ, while Y*^(0)^ *only captures the effect of the ancestral population through pure drift*.

*Then Y* ^(ℒ)^ *has expectation* **0** *and variances* **Φ**^(ℒ)^ *and* **Ω**^(ℒ)^; *Y* ^(0)^ *has expectation m*_0_**1** *and variances* **1** *and* **Ω**^(0)^; *and we have:*

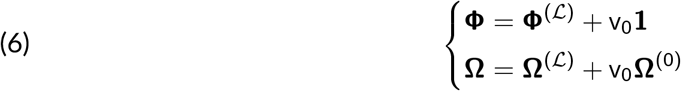

*with* **1** *the vector of ones. At the root* 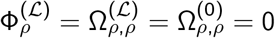. *Then we have by preorder recursion:*

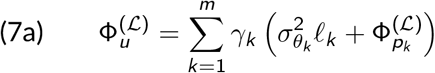

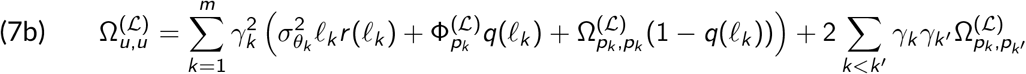

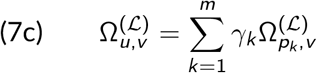

*and*

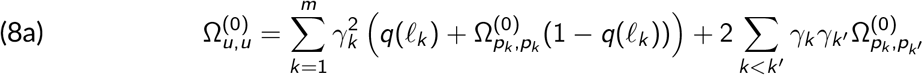

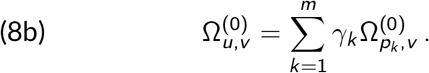

*Proof*. The additivity and recursion formulas directly stem from Equations (5a)-(5c) of Prop. 2. □

This decomposition means that we can separate the evolution process of Def. 3 into the two independent contributions of the trait evolution *Y* ^(ℒ)^ and of the random drift *Y* ^(0)^. Also, (8) clarifies that v_0_ impacts the model via Ω^(0)^, which solely depends on the network’s branch lengths in coalescent units.

#### 2.3.2. Generic branch length transformation

In traditional PCMs, models such as the Ornstein-Uhlenbeck or the Early Burst are commonly fitted using an equivalent BM on a tree with modified branch lengths (Ho and Ané, 2014). In this section, we show that our process on a network can also be re-written as the sum of two Brownian motions on modified phylogenies. Although the general formulas derived here have no direct computational use, they are used in Section 2.4 to link the model to a population genetics framework, and in Section 3.1 for an efficient implementation when the phylogeny is a tree.

Ignoring ILS, the Brownian motion with constant variance rate *σ*^2^ can be used along a species network (e.g., Pickrell and Pritchard, 2012). Under the BM model, the vector of traits at the tips is Gaussian, with a variance-covariance matrix *σ*^2^**C** that can be computed by traversing the tree in a preorder as (Bastide et al., 2018):

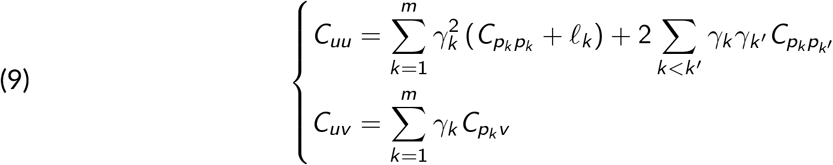

for any node *u* with parent nodes *p*_*k*_, parental branches of length *ℓ*_*k*_ and inheritance probabilities *γ*_*k*_ (1 ≤ *k* ≤ *m*), and *v* any node that is above *u* in the preorder. These propagation equations bear similarities with the equations of Cor. 4. Indeed, we can see the variance matrices **Ω**^(ℒ)^ and **Ω**^(0)^ as formally emerging from two different Brownian motions on the network, with rescaled branch lengths (see SM Section A for the proof):

##### Proposition 5

(BM on a rescaled network). *The variance matrix* v_0_**Ω**^(0)^ *on network N is the same as under the BM with variance rate* v_0_ *on a network with N ‘s topology and rescaled branch lengths:*

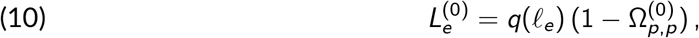

*for edge e with parent node p. Similarly, assuming a constant rate* 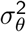 *across branches for simplicity, so that the scaled covariances and variances* 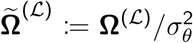 *and* 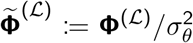 *do not depend on* 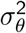, *then* **Ω**^(ℒ)^ *is the same covariance structure as under a BM with variance rate* 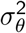 *and rescaled branch lengths:*

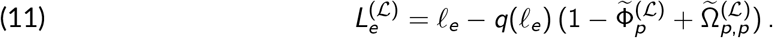

*Further, both transformations* (10) *and* (11) *are additive: for any nodes u* → *v* → *w along one lineage such that v is a tree node, then L*_*uw*_ = *L*_*uv*_ + *L*_*vw*_, *where L*_*uw*_ *is the transformed branch length of the combined edge going from u to w*.

The computation of the rescaled branch lengths involves one preorder traversal of the tree, but these lengths do not depend on the parameters 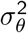 or v_0_. This transformation cannot be used in general as a computation technique (except in some special cases explored below), but ensures that the structure of the covariance matrix can be framed as emerging from a BM on the phylogeny. Note that the constant rate assumption is only technical. It leads to rescaled branch lengths 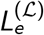 (and 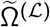) that do not depend on 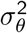. In SM Section A (Prop. S3), we describe a branch length transformation that depends on the rates 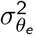 but can still cast **Ω**^(ℒ)^ as emerging from a BM with varying rates on the phylogeny.

#### 2.3.3. Special case of a phylogenetic tree

When the phylogeny is a tree, the recursion formulas and branch length transformations lead to closed-form expressions.

##### Proposition 6

(Closed-form expression in the tree case). *When the phylogeny is a tree, using the same model and notations as in Cor. 4, we get, for any two nodes u and v:*

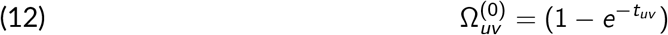

*where t*_*uv*_ *is the length of the unique path between the root and the MRCA of u and v (in coalescent units). Using* (10), *this leads to the following explicit branch length transformation:*

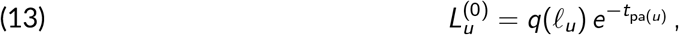

*where ℓ*_*u*_ *and* 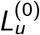 *are the original and transformed lengths of the branch going into u, and t* _pa(*u*)_ ^*is*^ *the length of the path between the root and the parent of u. Similarly, assuming for convenience a constant rate* 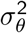 *on all branches:*

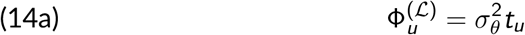

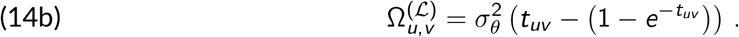

*Using* (11), *this leads to the following explicit branch length transformation:*

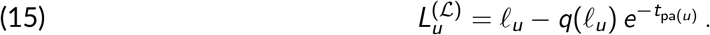

The proof is presented in SM Section A. Note that both transforms yield positive lengths, as 0 ≤ 1 − *e*^−*x*^ ≤ *x* for any *x* ≥ 0.

### 2.4. Phylogenetic model for the mean trait in populations^∗^

This section explores links between our model and established models in population genetics, e.g. for allelic frequencies. It can be skipped to focus on the practical use of the model.

#### 2.4.1. Mean population trait evolution on a network

The previous section tracks the trait expectation and variance in individuals sampled from each population. Another way to describe the model is to track the evolution of the trait distribution in a population, summarized by its mean (across all individuals in the population) and its within-population variance.

More precisely, define *M*_*u*_ as the average trait among all individuals in population *u* for a given locus and *V*_*u*_ its variance among individuals in *u*, that is, *V*_*u*_ is the within-population variance. Also define the expected within-population variance as *H*_*u*_ = E [*V*_*u*_] at node *u*. Then the moments of *Y* are linked to those of *M*_*u*_ and *V*_*u*_ by the following formulas.

##### Lemma 7

(Population mean and variance). *Write* **M** *and* **V** *the vectors of trait population means and variances at all nodes as above. Then:*

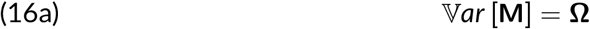

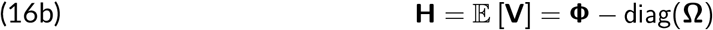

*where* diag(**Ω**) *is the vector made of the diagonal elements of* **Ω**.

The proof of this lemma is in SM Section A. By (16a), the covariance ℂov [*M*_*u*_; *M*_*v*_] between the trait means of two populations is equal to the covariance Ω_*u,v*_ between two different individuals randomly drawn from these populations. From (16b) we see that Φ_*u*_, the trait variance of an individual from a population *u*, is the sum of the covariance Ω_*u,u*_ between two individuals from *u* plus the expected within-population variance of the trait *H*_*u*_.

#### 2.4.2. Drift model for allele frequencies

A special case of interest is when *Y* represents the genotype of an individual at a given locus, with *Y* = 0 if the individual has the major allele, and 1 otherwise. In general, a classical Markov model of evolution for *Y* is not a Lévy process, as the increments are not independent. However, the special case with no mutations along the network can be represented with our model, setting 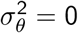. This special case corresponds to the “infinite site assumption” by which any new mutation affects a new locus so each locus displays at most one mutated allele, and further assuming that all mutations occurred prior to the root population *ρ*. Then, using notations from Cor. 4, *Y* = *Y* ^(0)^ from random drift due to ILS. The minor allele frequency *p*_*u*_ in population *u* is then the average of *Y* over all individuals in *u*, which is exactly *M*_*u*_. As *Y* can only be 0 or 1, *Y* ^2^ = *Y* and the population variance has a simple expression:

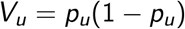

so that 2*V*_*u*_ is equal to the heterozygosity in population *u*. In that case, the expectation and variance in the root population are *m*_0_ = *p*_0_ and v_0_ = *p*_0_(1 − *p*_0_), and the covariance matrix between allele frequencies at all nodes is given by the matrix *p*_0_(1 − *p*_0_)**Ω**^(0)^, as computed from Cor. 4. The total variance **Φ** is constant equal to *p*_0_(1−*p*_0_), while the expected half heterozygosity is 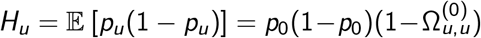. The expected *f*_2_ statistic used in population genetics can then be expressed, on a tree edge *e* going from *v* to *u*, as:

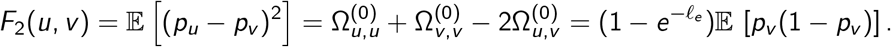

We recover the formulas from Patterson et al. (2012) when conditioning on *p*_*v*_ . If the phylogeny is a tree, **Ω**^(0)^ is also identical to the covariance used in Mary-Huard and Balding (2023).

These equations correspond to the population genetics framework that aims at recovering admixture graphs from frequencies of biallelic markers, using tools such as `TreeMix` (Pickrell and Pritchard, 2012) or `admixtools` (Maier et al., 2023; Patterson et al., 2012). In particular, if we assume that the minor allele frequencies (*p*_*u*_)_*u*_ are jointly Gaussian, and that branch lengths in coalescent units are small so that 1 − *q*(*ℓ*) ≈ 1 for all branches, then the propagation equations from Cor. 4 for the covariance become:

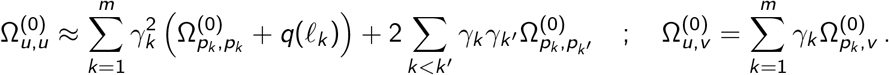

We recognize then the propagation formulas derived in Pickrell and Pritchard (2012) and used in many statistical tools such as `poolfstat` (Gautier et al., 2022) or `AdmixtureBayes` (Nielsen et al., 2023). These formulas also match (9), so that using this approximation, the model for the allele frequency is equivalent to a BM on the network, with branch lengths rescaled as *q*(*ℓ*) = 1 − *e*^−*ℓ*^ . As noted in Patterson et al. (2012), this rescaling of branch lengths is not additive: taking tree nodes *u* → *v* → *w* along one lineage, with *ℓ*_*uw*_ = *ℓ*_*uv*_ + *ℓ*_*vw*_, the rescaled *q*(*ℓ*_*uw*_) = 1 − 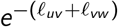 is *different* from *q*(*ℓ*_*uv*_) + *q*(*ℓ*_*vw*_) = 1 − 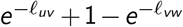. This is because the approximation ignores the fact that the drift decreases as we get away from the root.

Prop. 5 and Prop. 6 provide an exact branch length transformation, as demonstrated in the following corollary.

##### Corollary 8

(Allele frequency evolution as a Brownian motion). *The variance-covariance structure of the vector of allele frequencies p*_*u*_ *on the network is the same as the one obtained from a Brownian motion with variance p*_0_(1 − *p*_0_) *on the network, with branch lengths rescaled to*

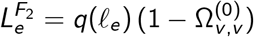

*for edge e, where v is the parent node of e. This transformation is additive, and transformed lengths can be seen as measuring “F*_2_*-units” (Gautier et al., 2022*; *Patterson et al., 2012)*.

*If the phylogeny is a tree, then* 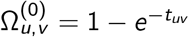 *and the transformation becomes explicit:*

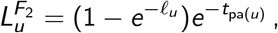

*writing* pa(*u*) *the unique parent node of u, and identifying an edge by its child node u. The back transformation from F*_2_*-units to coalescent units is then:*

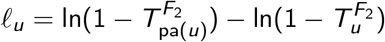

*with* 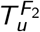 *the transformed length of the path from the root to u*.

The classical approximation from Pickrell and Pritchard (2012) then amounts to assuming constant expected heterozygosity along the phylogeny, that is, setting 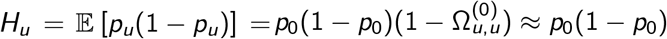. This leads to 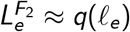 in Cor. 8.

### 2.5. The Gaussian-Coalescent model

We summarize here the core GC model for a univariate trait. The computations exploit the recursion formulas of Section 2.2 on a generic phylogenetic network, and we further derive a closed form for the GC variance matrix on a phylogenetic tree. In practice, we are interested in the distribution of the trait **X** = (*X*_1_, …, *X*_*n*_) measured on *n* individuals sampled from the populations at the leaves of a phylogeny, which may be a tree or a network with reticulations. In many applications, only one individual is sampled from each population but we do not make this assumption here.

#### Gaussian limit and the Gaussian-Coalescent model

In this section, we assume that *X* is a polygenic trait evolving according to Def. 4. Prop. 2 allows us to specify the expectation and variance for the vector **X** of measured traits.

##### Corollary 9

(Covariance of sampled traits). *Let* **X** = (*X*_1_, …, *X*_*n*_) *be the trait measured on individuals sampled from the tips of a species phylogeny. Assume that there are n*_*s*_ *different populations, and that n*_ind,*u*_ *individuals are sampled from population u, with* 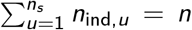. *We re-write this vector as* 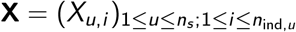, *where X*_*u,i*_ *denotes the trait for individual i in population u. Then, for any two populations u and v and distinct individuals i and j, we have:*

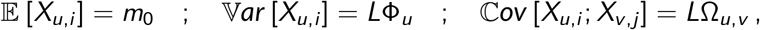

*where* **Φ** *and* **Ω** *are computed from Prop. 2. Using* **H** = **Φ** − diag(**Ω**) *as in* (16), *the variance of* **X** *is the sum of the between-population variance* **Ω** *and the expected within-population variance* **H**:

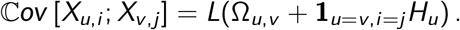

In general, the distribution of **X** is not Gaussian. However, from the central limit theorem, the vector of traits **X** can be approximated by a Gaussian vector when *L* is large.

##### Proposition 10

(Gaussian limit). *When the number of loci L goes to infinity, the trait vector* **X**, *with elements* 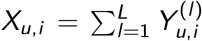 *across nodes* 1 ≤ *u* ≤ *n*_*s*_ *and individuals* 1 ≤ *i* ≤ *n*_ind,*u*_, *converges in distribution towards a Gaussian vector* **Z**.

This proposition is a direct consequence of the multivariate central limit theorem, as the polygenic trait is the sum of independent identically distributed random vectors **Y**^(*l*)^ with finite variance matrix. It allows us to fit a statistical model to a vector of observed trait **X**, using the Gaussian approximation. In all statistical analyses below, we use the “Gaussian-Coalescent” model, which approximates the coalescent-based process by a Gaussian vector with an expectation and a covariance matrix as in Prop. 10. This joint Gaussian approximation was already used in previous work (Hibbins et al., 2023; Mendes et al., 2018). We note that it remains an approximation even if all loci follow a Brownian motion, as demonstrated in SM Section A (Remark S1) and illustrated in SM Section B.

#### 2.5.2. Closed-form expression for species trees

In the special case when the species phylogeny *N* is a tree, a direct consequence of Cor. 4 and Prop. 6 is the analytical formulation of the covariance matrix, beyond the recursion formula, in the following proposition. We use it later to compare our model with ILS to other models, without ILS or from alternate approaches taken by other authors.

##### Proposition 11

*Assume a phylogenetic polygenic evolution model on a species tree with a finite and constant variance rate (per coalescent unit)* 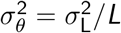 *for the Lévy distribution of increments at each locus, and trait distribution P*_0_ *at the root ρ with mean m*_0_ *and variance* v_0_. *If t*_*u*_ *and t*_*uv*_ *denote the path and shared path lengths in coalescent units, then the conditional trait means and conditional (co)variances between individuals from population(s) u and v, conditional on ρ, are* E [*X*_*u,i*_] = *m*_0_ *and*

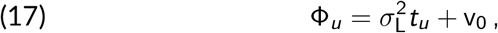

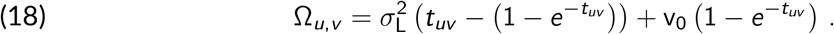

*Writing* 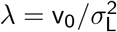 *as in* (1), *we get:*

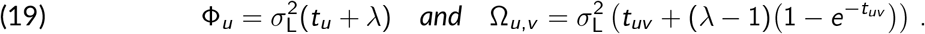

##### Remark 2

(interpretation). Using notations from (2), we can rewrite (18) as:

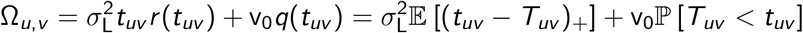

where *T*_*uv*_ is the random coalescent time between two individuals, one from population *u* and the other from population *v*, that are allowed to coalesce only starting from the branch above their most recent common ancestor. In Ω_*u,v*_, the first term is the expected shared time of evolution since the root scaled by the evolutionary rate 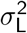, and the second term is the ancestral variance v_0_ times the probability that they have coalesced during the allowed time *t*_*uv*_ . Up to a constant, this formula has the same general form as eq. (11) from Schraiber et al. (2024), who assumed a compound Poisson model of evolution on *L* loci and v_0_ = 0.

##### Remark 3

(within-population variance). The expected within-population variance of the trait is

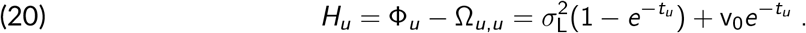

It is equal to the equilibrium variance 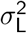 when *λ* = 1, and otherwise tends to it when *t*_*u*_ goes to infinity, that is, for a deep tree, or a tree with a root being pushed back in time with a single long branch. Our model hence recovers the expected equilibrium variance (Lynch and Hill, 1986).

##### Remark 4

(root variance at equilibrium). When we assume that the root variance is at equilibrium, i.e. 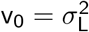 or *λ* = 1, then (19) and (20) give:

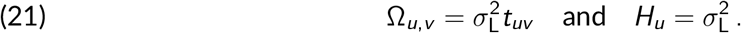

The process is then the same as the standard BM on the tree in coalescent units, but with added within-population variation, with a variance at each individual increased by one coalescent unit compared to the covariance between individuals from the same population.

#### 2.5.3. Equivalence to the BM with transformed branch lengths

In this section we continue to assume that a *N* is a tree. We can see present-day populations as internal nodes, with extra branches leading to each sampled individual, initially of length 0 if there is no within-species variation. The covariance of our model with ILS is then equivalent to that of a standard BM with variance 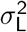 on this extended tree, with length *ℓ*_*u*_(*λ*) for the internal edge to population *u* and *ℓ*_*u,i*_ (*λ*) for the external edge to individual *i* sampled from *u*, given by

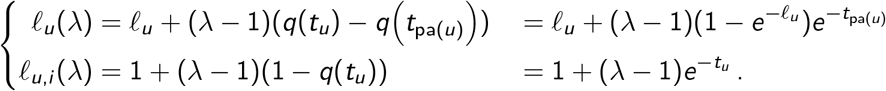

For internal branches, the transformation directly stems from (13) and (15) of Prop. 6. These give the covariance structure between different individuals, that exactly matches **Ω**. For external branches, the transformation accounts for the within-population variation **H** = **Φ** − diag(**Ω**), and stems from (14) of Prop. 6.

If *λ* = 1 (root at equilibrium), then the internal branch lengths are unchanged, and the tip branches are assigned one coalescent unit: the process reduces to a standard BM with within-population variation. Increasing or decreasing *λ* leads to an increase or decrease of all branch lengths.

This equivalence to a standard BM allows us to use the three-point structure algorithm of Ho and Ané (2014). Like for other models such as the Ornstein-Uhlenbeck or the Early Burst, this tree transformation can be combined with an extra non-heritable independent within-species variation by lengthening the external branches, as implemented in `phylolm` (Ho and Ané, 2014).

#### 2.5.4. Population size as inverse phylogenetic signal

In the case when *N* is a tree as earlier, and assuming a constant haploid effective population size 2*N* over *N*, we study in SM Section C the limits in very small and very large populations. Under no ILS (*N* → 0 and immediate coalescence) the model is equivalent to a BM model on the original tree; while under maximum ILS (*N* → ∞and all coalescences above *ρ*) the model is equivalent to a single population (or star tree) with independent individuals. In short, the population size *N* can be seen as an inverse measure of “phylogenetic signal”: the larger it is, the weaker the correlations between populations (see also Mendes et al. 2018).

If branch lengths were known in number of generations, then we could imagine fitting *N* to the continuous traits at hand, so that our model would adapt to the specific level of coalescence affecting the genes controlling the traits.

### 2.6. Comparison with alternate approaches accounting for ILS^∗^

In this section, we explore similarities and differences with previous approaches that account for ILS using the theoretical distribution of gene trees from the coalescent. We did not consider methods that use an empirical distribution of gene trees as input, instead of a single species phylogeny. This section can be skipped to focus on the use of the GC model.

#### 2.6.1. The C^∗^ matrix

Mendes et al. (2018) proposed the first approach accounting for ILS, in the tree case. As here, they considered a phenotype controlled by many independent and additive loci. They assumed an ultrametric tree (with all taxa at the same distance from the root), and one individual per taxon. Their trait covariance is obtained by averaging the covariance matrices associated with gene trees, weighted by their probability under the coalescent process. For each gene tree, the associated matrix corresponds to the covariance conditional on the ancestral value at the root of the tree, that is, at the most ancient coalescent event between the sampled individuals. When deriving the BM covariance on a tree, this root value is taken as known (fixed to 0 in Mendes et al., 2018, S.13), or as a parameter to be estimated. Each gene tree covariance then corresponds to conditioning on the locus effect at a random time, and these times differ across loci. In contrast, our model conditions on a fixed ancestral population, in which ancestral polymorphism is modeled via the distribution *P*_0_.

Mendes et al. (2018) found the following general expression for their covariance with ILS, in their equations (S.14) for the variance case *u* = *v* and (S.17) for the covariance case *u*≠ *v* :

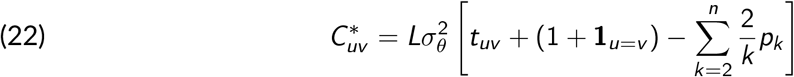

where *n* is the number of populations (tips) in the species tree, 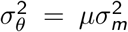 is the compound Poisson model with rate *µ* and mutation variance 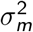, and *p*_*k*_ is the probability that *k* lineages of the gene tree enter the population of the MRCA of all *n* populations (going back in time). For example, if the tree has *n* = 3 populations and an internal edge length of *t* coalescent units, then *p*_2_ = (1 − *e*^−*t*^) and *p*_3_ = *e*^−*t*^ . In general, *p*_*k*_ depends on the tree size (number of sampled populations *n*), topology and edge lengths in coalescent units. When 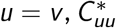 is the phenotype variance of one individual in population *u*, similar to Φ_*u*_ in Prop. 11.

##### Remark 5

(impact of taxon sampling). In the model derived by Mendes et al. (2018), the covariance between two populations depends on all other populations sampled in the tree, to know the MRCA of the entire sample in each gene tree. This means that adding or removing a tip somewhere in the tree impacts the covariances of all the other populations (see Sec. 2.6.2 below). This is not the case with the GC model provided that the root population is not affected by the taxon addition or removal (as e.g., in Figure 2), as can be seen directly in Prop. 11 on a tree, and by Prop. 3 generally. The model derived by conditioning on a given root *ρ* enjoys the following property: if *S*_1_ is a “large” sample of individuals and populations, and if *S*_2_ is a subset of individuals and/or populations from *S*_1_, then the model defined for 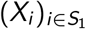 on the full *S*_1_ reduced to data on *S*_2_ is equivalent to the model defined directly for 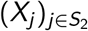 on *S*_2_ only.

**Figure 2.**
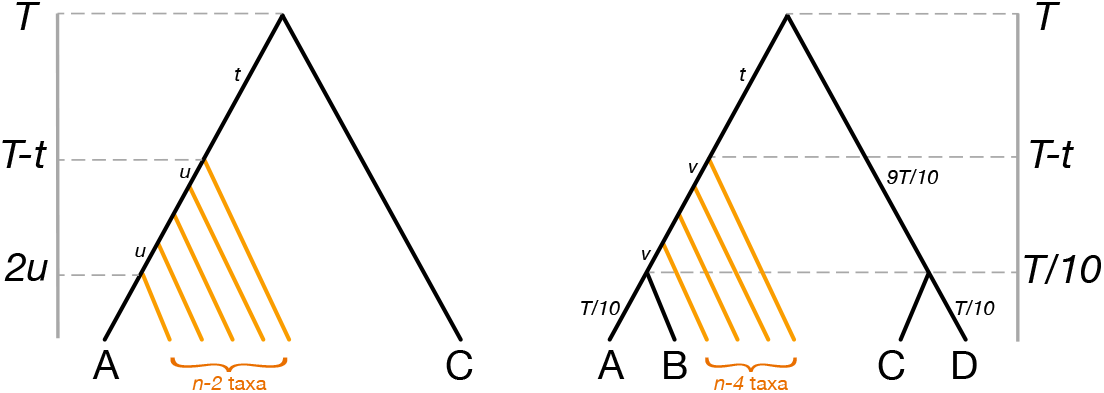
Species trees used to quantify the effect of taxon sampling on ILS-based methods, each having *n* taxa at distance *T* from the root. The gray axis indicates time before present (from 0 at the tips to *T* at the root). Branches are annotated by their length. **Left**: tree used in Fig. 3, with *n* − 2 unlabeled taxa (shown with orange branches) forming a clade with *A*. This clade’s MRCA is at distance *t* from the root, with successive speciation events separated by branches of length *u* = (*T* − *t*)*/*(*n* − 1), such that A is at distance *T* from the root. **Right**: tree used in Fig. 4, with *n* 4 unlabelled taxa (orange) forming a clade with *A* and *B*. This clade is at distance *t* from the root, with successive speciation events separated by branches of length *v* = (9*T /*10 − *t*)*/*(*n* − 4), such that all species are at distance *T* from the root, and the MRCA of A & B is at distance 9*T/*10 from the root.

##### Remark 6

(3-taxon tree with root at equilibrium). On a 3-taxon species tree with an internal branch length of *t* coalescent units, (22) simplifies to:

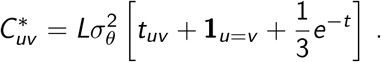

Taking *λ* = 1 in (19), the GC model leads to

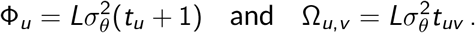

When there is only one individual per population, the variance matrix of the GC model is then:

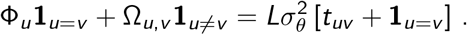

In this specific case, the two formulas agree up to a uniform term 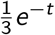, that goes to zero when *t* goes to infinity. Note that the actual value of this term does not matter to estimate parameter values: both covariance matrices lead to the same estimated ancestral state at the root and the same variance rate (but differ in their estimated standard errors). Our examples below show that this agreement between *C*^∗^ and our GC model covariance does not generalize to more than 3 taxa.

#### 2.6.2. Impact of taxon sampling: numerical experiments

Hibbins et al. (2023), who coined the name ‘*C*^∗^’, extended the approach of Mendes et al. (2018), which takes *C*^∗^ as the average of covariance matrices across gene trees, and implemented it in the R package `seastaR`. They consider gene trees generated from the coalescent on a species tree (as in Mendes et al., 2018) or observed from a sample of loci. Their coalescent-based species-tree method matches that of Mendes et al. (2018) on 3 taxa, but differs on 4 or more taxa, as they break down the full tree into 3-taxon subtrees and then calculate *C*^∗^ from (22) on each triplet.

To compare with the GC model in the tree case, we considered the phylogenies in Fig. 2 with focal populations *A* and *C*, and *n* sampled populations below their MRCA. For each phylogeny, we calculated the coalescent-based *C*^∗^ matrix implemented in `seastaR` (function `get_full_matrix`, Hibbins et al., 2023), and the GC covariance matrix (Prop. 11) assuming equilibrium variance in the root population (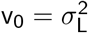, based on Remark 6). For both, we assumed one individual per taxon (as required by *C*^∗^) and calculated the *scaled* covariance matrix, omitting the variance rate factor 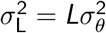.

We first considered the trait variance, at *A* and *C* in the phylogeny in Fig. 2 (left) in which *C* is sister to a clade containing *A*. Fig. 3 (left) shows these variances as a function of the clade’s age and size. One would expect Var [*X*_*A*_] and Var [*X*_*C*_] to be equal, because *A* and *C* are at the same distance from the root (fixed to *T* = 30 or 3). Also, one would expect these variances to be independent of the clade size and age, to avoid a dependence on taxon sampling in this clade. These expectations hold for the GC model but not for *C*^∗^. When the full phylogeny has *n* = 3 taxa (*A*, its sister, and *C*) then 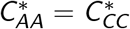 holds, but its value depends on the divergence time between *A* and its sister taxon (very slightly when *T* = 30 and less slightly when *T* = 3 coalescent units) — in agreement with Remark 6. However, when *A* is in a larger clade (*n* ≥ 4) then *C*^∗^ assigns a significantly smaller variance to *A* than to 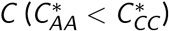, with a younger clade age (larger *t* in Fig. 3) leading to a larger gap.

**Figure 3.**
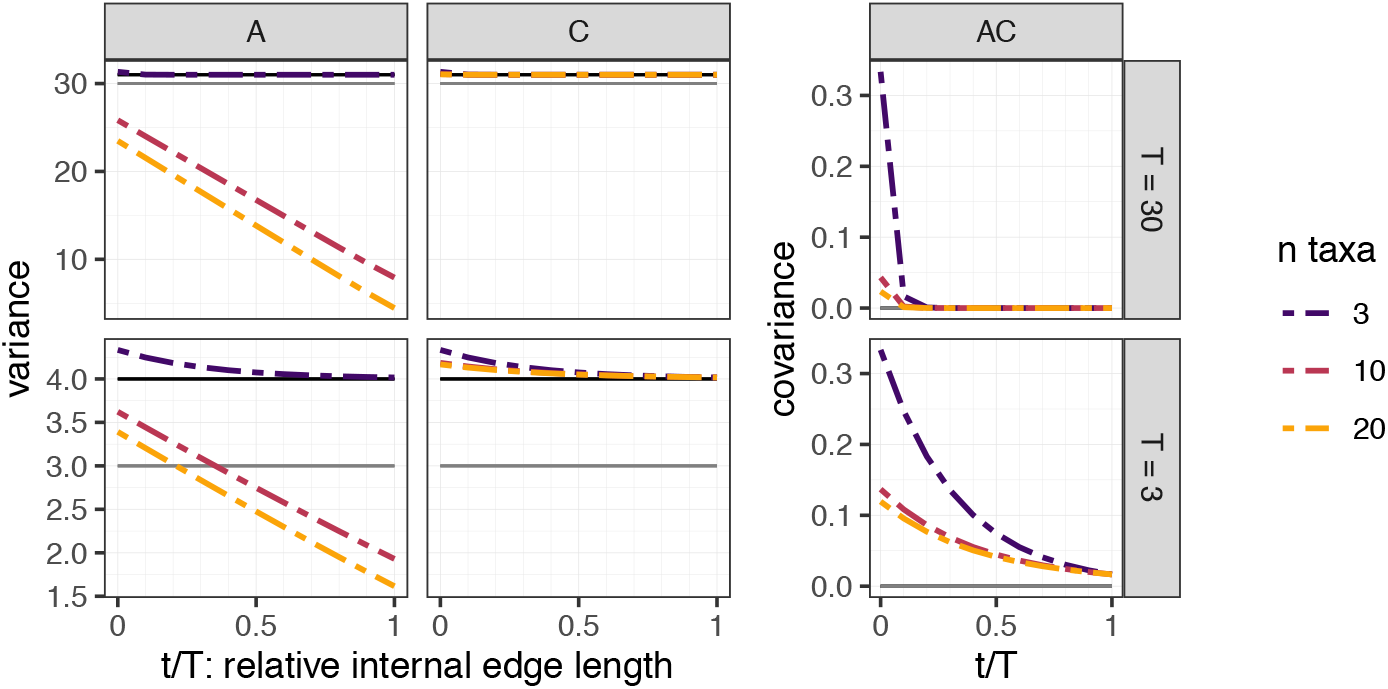
Effect of taxon sampling and clade’s distance to the root on the variance and covariance calculated by the ILS-based *C*^∗^ (Hibbins et al., 2023; Mendes et al., 2018), on the species tree in Fig. 2 (left) with branch lengths (*T* and *t*) in coalescent units. **Left**: trait variance at *A* and at *C* . **Right**: covariance between the traits at *A* and *C* . Colored dashed lines show *C*^∗^ values. The value from the GC model is shown by a black horizontal line (at *T* + 1 = 31 or 4 for the variance, 0 for the covariance). The value from the standard BM model (without ILS) is indicated with a dark gray horizontal line (at *T* = 30 or 3 for the variance, 0 for the covariance).

Intuitively, this is because *A* is part of triplets *AB*_1_|*B*_2_ where *B*_1_ and *B*_2_ are orange taxa in Fig. 2, whose MRCA is younger than the species tree root. In a gene tree, denote the age of the oldest coalescent event as 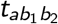 between *A, B*_1_ and *B*_2_, and 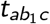 between *A, B*_1_ and *C* . Then 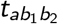 is typically younger: 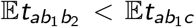. Conditional on the trait at the gene tree root, the variance of *A* calculated from the gene tree with *B*_1_ and *B*_2_ is only 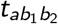, which excludes the variation in the trait between 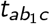 and 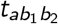. Ignoring this ancestral variation by conditioning on the trait value at a more recent time causes a reduction in 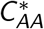. This reduction does not affect 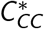, because *C* is not part of triplets whose MRCA is younger than the root of the species tree.

We then turned to phylogenetic covariances. Using the tree in Fig. 2 (left), ℂov(*X*_*A*_, *X*_*C*_) does not depend on taxon sampling in the GC model, with a covariance of 0 because we condition on the MRCA population of *A* and *C* (rather than a more ancestral population). Based on *C*^∗^, the phylogenetic covariance between *A* and *C* depends on taxon sampling, especially when the total tree height is low (*T* = 3 coalescent units) or when taxon sampling adds speciations near the root (*t* small, Fig. 3).

In Fig. 2 (right), we considered a phylogeny with 2 similar sister pairs: *AB* and *CD*. As both pairs share the same divergence time, one would expect the covariances ℂov(*X*_*A*_, *X*_*B*_) and ℂov(*X*_*C*_, *X*_*D*_) to be equal and independent of taxon sampling within the species tree. This expectation holds for the GC model but not for *C*^∗^ (Fig. 4). When no other taxa are sampled than the 2 sister pairs (*n* = 4) then 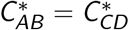 holds. But when *A* and *B* are part of a larger clade (orange in Fig. 2) then 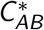 is significantly smaller than 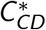, with a larger gap when the clade is younger (larger *t*).

**Figure 4.**
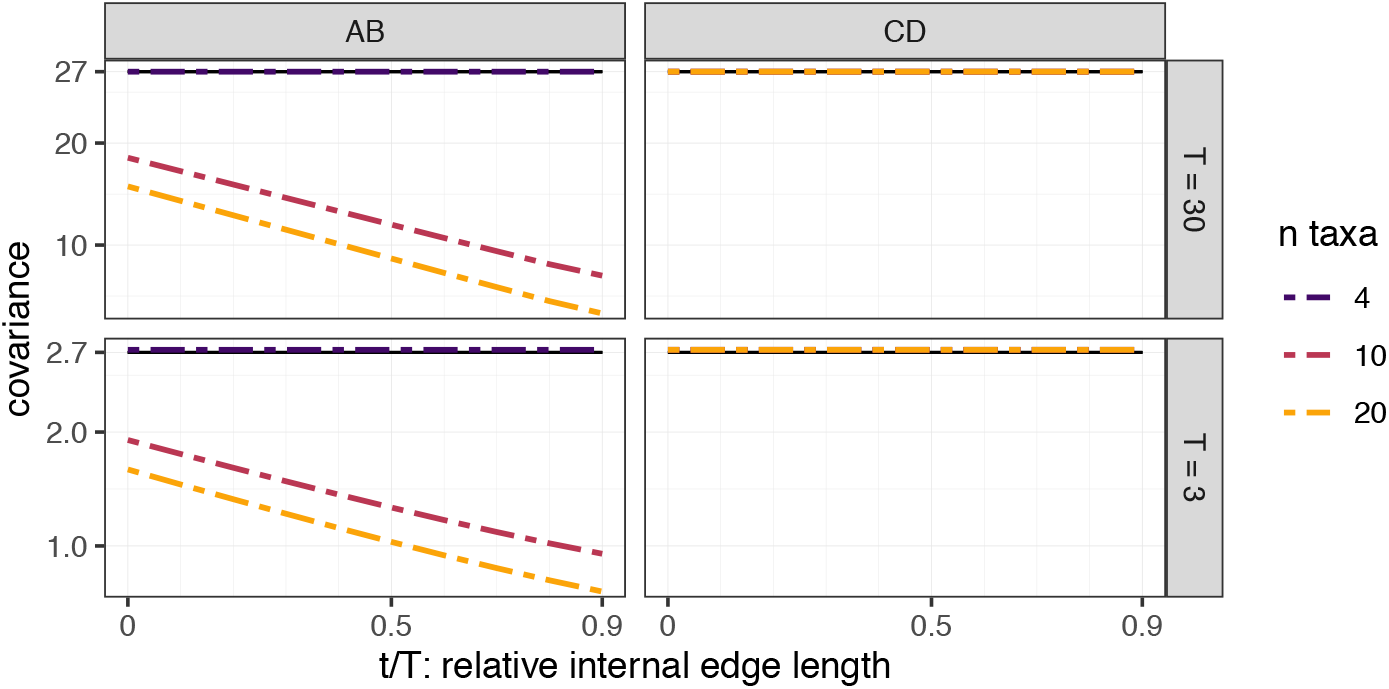
Effect of taxon sampling and clade’s distance to the root on the covariance *C*^∗^ of two sister pairs, on the species tree in Fig. 2, right, with branch lengths (*T* and *t*) in coalescent units. **Left**: trait covariance between *A* and *B*. **Right**: trait covariance between *C* and *D*. The value from the GC model is shown by a black horizontal line. It is identical to the value from the standard BM model (without ILS), 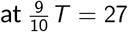 (top) or 2.7 (bottom).

Intuitively, this is because the *AB* pair is part of triplets *AB*|*B*_2_ whose MRCA is younger than the species tree root, whereas all triplets containing the *CD* pair have their MRCA at the root. By conditioning on the trait at the younger root of gene trees, a shorter evolutionary path is considered to be shared between *A* and *B*, causing a reduction of their covariance compared to the *CD* pair.

Averaging over all triplets to get *C*^∗^ could be partly responsible for some of the sensitivity of *C*^∗^ to taxon sampling. To reduce this sensitivity, a future version of `seastaR` may use a subset of triplets only, such as those spanning the root of the tree (an idea by M. Hahn and M. Hibbins in their reviews, see Rogers 2026). We used here the released version of the software at the time of the study (`v0.1.0#9c6cf78`), which is also the version used in Hibbins et al. (2023).

Another avenue to reduce this sensitivity is to follow a procedure by Mo and Hahn (2026): simulate many gene trees from the species phylogeny, then average their associated BM co-variance matrices using function `trees_to_vcv` in `seastaR`. This alternative *C*^∗^ estimate is still sensitive to taxon sampling, and sometimes more so than `get_full_matrix` (see appendix D.1).

#### 2.6.3. Introgression and ILS

To account for introgression in addition to ILS, Hibbins and Hahn (2021) extended the *C*^∗^ method in the specific case of 3 populations, assuming one bidirectional gene flow event between the outgroup population and one ingroup population. They obtained the phylogenetic trait covariance as a weighted average of *C*^∗^ derived from 3 parent species trees: one representing no introgression, and two representing unidirectional introgression. Our polygenic model has the advantage of being applicable to any phylogenetic network on any number of populations and individuals, following the multispecies network coalescent exactly, and with the guarantee that given a fixed root population *ρ*, variances and covariances are independent of taxon sampling.

## 3. Impact on comparative methods

### 3.1. Phylogenetic regression and implementation

We implemented a phylogenetic regression whose residuals are Gaussian with a variance matrix structured as the GC, assuming a constant rate across branches. The regression takes as input: (i) a vector of univariate continuous traits **X** = (*X*_1_, …, *X*_*n*_) measured on *n* individuals sampled from the populations at the leaves of a phylogeny, with possibly several measured individuals per population; (ii) a phylogeny with branch length in coalescent units and (iii) a regression design matrix **R** with *n* rows and columns that may include an intercept (vector of ones), structured vectors for phylogenetic ANOVA, and/or covariates of interest. The phylogenetic regression models **X** using the covariance from the GC model in Cor. 9, denoted 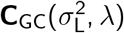 here:

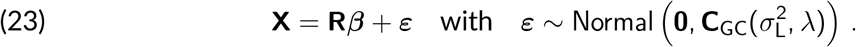

This covariance can be computed from the recursion in Prop. 2, or using the closed form from Prop. 11 for a tree. The model parameters are the vector of coefficients ***β***, the rate 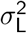 per coalescent unit, and the ratio 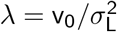 between the root variance and the rate.

For example, **X** may contain a measure of the constitutive defense response to herbivory, observed across multiple plants from various species and populations. If the design matrix **R** is reduced to a vector of ones, then ***β*** has only one coefficient (the intercept) *β*_1_ = *m*_0_, that represents the mean response in the ancestral population. To test for a negative correlation between constitutive defense and extra defense induced by prior herbivory, **R** may be extended with a second column containing induced defense values (as in Haak et al., 2014), with associated regression coefficient *β*_2_ and *β*_2_ *<* 0 providing evidence for a negative relationship between the two defense types. (See e.g., Harmon 2019 for other application examples of phylogenetic regression.) The design matrix **R** can also be constructed to allow for different mean values among different groups of populations, as in a typical ANOVA. This amounts to allowing for shifts in the trait mean, as is done, e.g., in Bastide et al. (2018) to test for transgressive evolution after hybridization events.

As in classical phylogenetic regressions (see e.g., Ho and Ané, 2014; Teo et al., 2023), the variance parameters are fitted using restricted maximum likelihood (REML) with a numerical optimization of *λ*, and ***β*** is estimated with maximum likelihood. Tests, confidence intervals and ancestral state reconstructions can then be performed conditionally on *λ* as in the classical setting (see e.g., Bastide et al., 2018). Similarly, classical penalized likelihood scores such as AIC can be used to compare this GC phylogenetic regression with other models. For example, one may compare the GC, BM and Ornstein-Uhlenbeck covariance models, or fit the GC model across multiple phylogenies with various reticulation events to test how introgression helps explain the trait data (Karimi et al., 2020; Morley et al., 2026; Teo et al., 2023). This model could also be used to infer the phylogeny from continuous traits, provided that data are combined across many independent traits. For instance, admixture graphs are commonly reconstructed from allele frequencies (e.g., Nielsen et al., 2023; Patterson et al., 2012; Pickrell and Pritchard, 2012), that can be modelled with the GC (see Section 2.4.2).

We implemented the GC model in the `julia` package `PhyloTraits` (Bastide et al., 2018) v.1.2.0, which can handle phylogenetic networks. We also implemented the GC model in the `R` package `phylolm` (Ho and Ané, 2014) v2.7.0, which is restricted to species trees, but can estimate non-heritable trait variation in addition to the within-species heritable variation expected from the GC model, as mentioned above. Both the `julia` and `R` implementations can also perform phylogenetic linear models (ANOVA and multiple regression) using predictors and the Gaussian-Coalescent model of phylogenetic correlation for residuals.

### 3.2. Estimation under ILS and gene flow

We performed simulations to quantify the impact of accounting for ILS using the GC model, versus ignoring ILS or using *C*^∗^ from the `seastaR` package (in the tree case). We looked at the accuracy of the estimated rate of trait evolution, as in Mendes et al. (2018) and Hibbins et al. (2023).

#### 3.2.1. Simulations settings

To simulate polygenic traits controlled by loci whose gene trees were generated under the coalescent, we used `PhyloCoalSimulations` (Fogg et al., 2023) v1.1.0. We generated each locus under a compound Poisson distribution with Laplace-distributed effects. Specifically, the number of mutations was generated from a Poisson process, and each mutational effect was drawn from a centered Laplace (symmetric exponential) distribution with shape 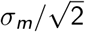, which has mutational variance 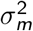. We used a large *L* = 100, from which the Gaussian approximation should be accurate.

For this simulation, we used the *Polemonium* phylogeny from Teo et al. (2022): a 17-taxon network with 3 reticulation events estimated in Rose (2021) and calibrated in Teo et al. (2023). From each taxon, we simulated *n*_*i*_ = 20 individuals. To simulate either low or high ILS, branch lengths in the phylogeny were rescaled to a total height of *T* = 10 (low ILS) or *T* = 1 (high ILS) coalescent units. Each locus effect had *µ* = 3*/T* mutations per coalescent unit, for an expected 3 mutations along gene trees from the root population to each individual. Mutation effects had mutational variance 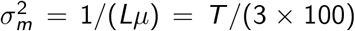, such that the trait variance rate was 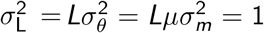. The ratio 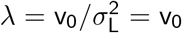 of the phenotype root variance over its equilibrium variance was varied in {0, 0.5, 1, 2}. As the phenotype’s root variance is *L* times that of the locus effect’s root variance, each locus was taken to have variance v_0_*/L* in the root population. The mean in the root population was fixed to *m*_0_ = 0. We simulated 100 independent replicates for each parameter combination. While 17 taxa is a small sample size to estimate the evolutionary variance rate, *n*_*i*_ = 20 individuals per population was expected to provide information on *λ*, or the ancestral trait variance v_0_ at the root.

For each data set, we used the `gaussiancoalescent` model in `PhyloTraits` v1.2.0 (Bastide et al., 2018) to estimate the evolutionary variance rate 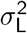, either assuming variance equilibrium at the root (*λ* = 1, which was not necessarily the true value), or assuming the true variance at the root, or simultaneously estimating *λ*. We also analyzed each data set under a Brownian motion model, to estimate the evolutionary variance rate 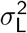 simultaneously with a within-population variance 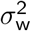, assumed shared across all populations. Specifically, this model assumes a BM with rate 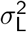 for the evolution of the population means, which are not observed. Conditional on these means, an observed individual value is assumed normally distributed centered at its population mean and with variance 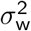. See Teo et al. (2023) for details. All the variances were estimated using the restricted maximum likelihood (REML) procedure, and we used a phylogenetic regression with a single intercept representing the root population mean value *m*_0_. Given the reticulate phylogeny and the multiple individuals per species, estimation with `seastaR` was not applicable.

#### 3.2.2. Simulation results

Under low ILS, the evolutionary variance rate 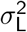 was estimated without practical bias regardless of the estimation model: under a BM, or under a GC with *λ* fixed or estimated (Fig. 5, top row). There was no noticeable gain to knowing the true value of *λ*. This is expected from theory for limit cases with no ILS (see Sections 2.5.4 and C). Further, precision increased greatly under any GC model compared the BM model. For example, the average root-mean-squared error (RMSE) on *σ*_L_ was 0.046 from the GC assuming *λ* = 1 versus 0.179 under the BM model. Under high ILS, 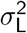 was estimated accurately only under the GC model with *λ* fixed to its true value (Fig. 5, bottom row). Without knowledge of *λ*, fixing *λ* = 1 was the best option, with error dominated by the bias (RMSE of 0.136). Bias was smaller when *λ* was estimated, but precision was lower. Under the BM model, bias was similar to the GC assuming *λ* = 1 but precision was lower, resulting in higher error overall (RMSE of 0.256). In short, the GC model fixing *λ* = 1 was the most accurate in all our simulation settings (in case *λ* is unknown).

**Figure 5.**
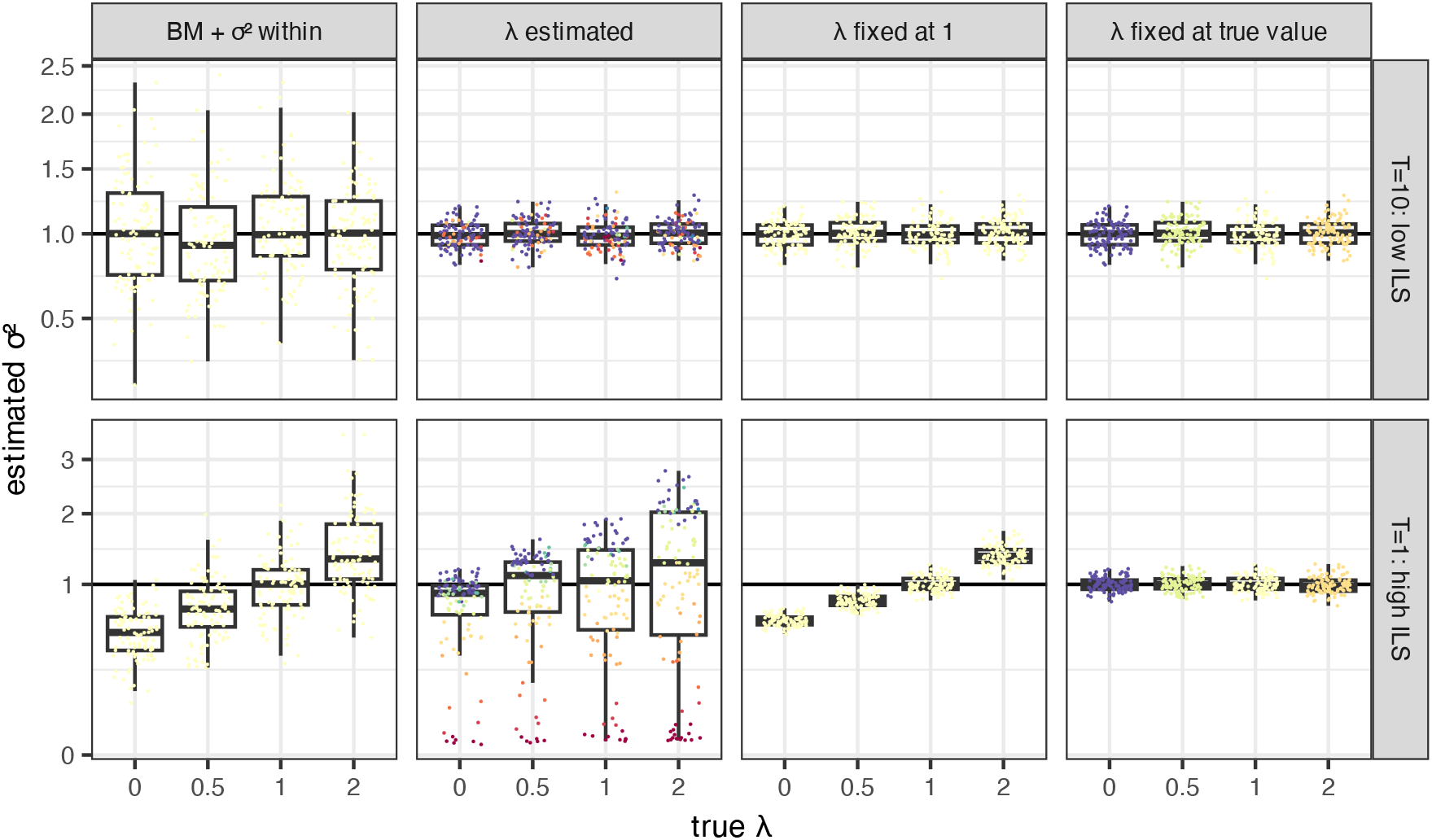
Estimated variance rate 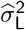 (displayed on the square-root scale) from data simulated under the *Polemonium* 17-taxon network, 20 individuals per taxon, and traits controlled by *L* = 100 loci with compound Poisson-Laplace effects. The true rate was 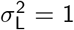. Top: low ILS with network height of *T* = 10 coalescent units. Bottom: high ILS, with *T* = 1. The top and bottom rows use different limits for the vertical axis. Each column corresponds to one estimation method: the BM model with extra within-population variation, the GC model with *λ* estimated, fixed at 1, or fixed at its true value (from left to right). Colors indicate the estimated or fixed *λ* (or yellow for the BM model), with cold colors for low values and warm colors for large values.

The ancestral state, or trait mean in the root population *m*_0_ was unbiased, from all estimation methods (Fig. S6 in SM Section D). Its precision was much better under high than low ILS. This is expected from theory again (see Section 2.5.4), as limit cases with maximum ILS correspond to a lack of phylogenetic correlation with independent individuals. Informally, gene genealogies trace back to more individuals from the root population at high ILS, leading to more information about the root population under high than low ILS. At low ILS, precision in *m*_0_ is driven by the number of populations and their phylogenetic correlation. At high ILS, precision in *m*_0_ is driven by the total number of individuals and the trait variance at the root, v_0_.

Under GC, the estimation of *λ* was extremely imprecise, especially under low ILS. Under high ILS, estimates of *λ* were slightly more concentrated around the true value (Fig. S7 in SM Section D). This is expected, as *λ* depends on the trait variance at the root, and because the trait data contain more information about the root population under high ILS.

Model selection also suggested that contemporary trait data carry little information about *λ*, at least in our simulation setting. Using a likelihood ratio test at level *α* = 0.05, *λ* = 1 was rejected in only 1.1% of replicates (black dots in Fig. S7). Using AIC, the GC model with fixed *λ* = 1 was favored 79% of the time, followed by the BM model 17% of the time. Estimating *λ* was deemed worthwhile only 4% of the time (Fig. 6).

**Figure 6.**
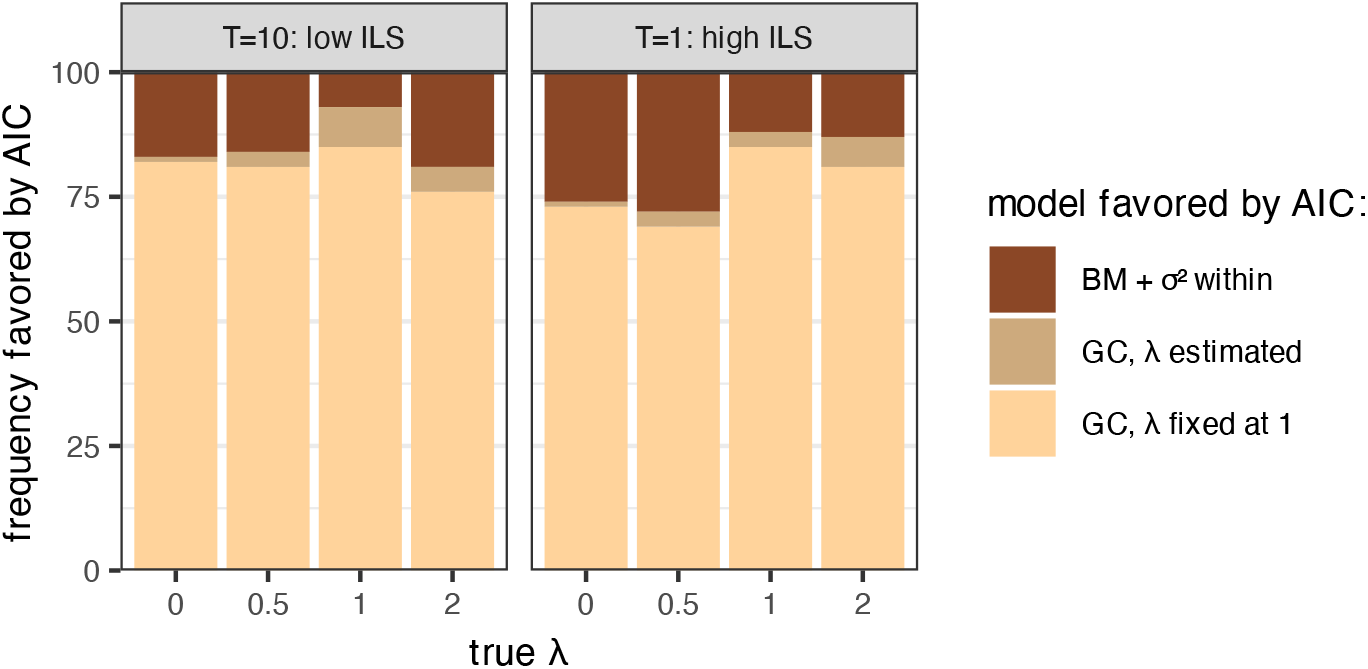
Proportion of times that each model was favored by AIC, out of 100 replicate data sets, when comparing 3 models: Brownian motion with intra-specific variation, Gaussian-Coalescent (GC) with *λ* estimated simultaneously with 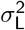, and GC with *λ* = 1 fixed.

The ancestral trait variance v_0_, like *λ*, was poorly estimated (Fig. S8), unless the GC model was used with *λ* fixed to its true value. However, the expected within-population variance *H*_*u*_ was estimated accurately, at both ILS levels. For a population *u* that is not below any reticulation, the expected variance within *u* is given by (20): it is close to 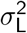 under low ILS and close to v_0_ under high ILS. For a population *u* whose history is affected by one or more reticulation, *H*_*u*_ can be calculated recursively. Intuitively, reticulation increases *H*_*u*_ (at fixed ILS, or fixed distance *T* between *u* and the root population), especially if admixture involved divergent parent populations. We calculated *H*_*u*_ for each population *u* based on the true model parameters, then considered the average within-population variance *h*, averaged across the 17 populations. For each simulated data set, we repeated this calculation of the average expected within-population variance, but using the estimated parameters, to obtain an estimate 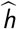 of *h*. When fitting with the BM model, we used the estimated within-species variance (assumed shared across populations) as 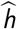. This within-population parameter *h* was well estimated at both ILS levels (Fig. S9). From 20 individuals per population, this is not surprising. However, considering that 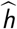 was calculated from very noisy estimates 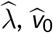, and 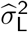 (noisy under high ILS) under GC, accurate estimation of 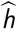 is still noteworthy.

Finally, we note that the “high ILS” setting was particularly challenging, as the entire network had a total height of only one coalescent unit. When reproducing this simulation study on the empirical tomato tree described below, which has a height of 6 coalescent units, results were in line with the “low ILS” setting above. In particular, the GC model with fixed *λ* = 1 received the most AIC support, and produced accurate estimation of 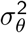, with an RMSE value of 0.120, much lower than the BM model (0.269), and almost equal to the best-case GC model with *λ* fixed to its true value (0.119). See SM Section D for details.

### 3.3. Floral traits in the wild tomato clade

#### 3.3.1. Methods

We re-analyzed the wild tomato dataset studied in Hibbins et al. (2023), that includes several floral traits (corolla diameter, anther length and stigma length) compiled in Haak et al. (2014) and a phylogenetic tree from Pease et al. (2016), dated in coalescent units assuming a population size of 100 000 and one generation every two years as in Hamlin et al. (2020).

We first reproduced the analysis of Hibbins et al. (2023), who used two different triplets extracted from the phylogeny (see Fig. 7): a “high ILS” triplet (tree height of 0.5 coalescent units) consisting of *S. galapagense* (LA0436), *S. cheesmaniae* (LA3124), and *S. pimpinellifolium* (LA1269); and a “low ILS” triplet (tree height of 6 coalescent units) with *S. pennellii* (LA3778), *S. pennellii* (LA0716), and *S. pimpinellifolium* (LA1589). IDs are accession numbers from the Tomato Genetics Resource Center. For each population (accession) and trait, we took the average over all the individual measurements, and then fitted 4 models to each triplet: the simple BM, the `seastaR` method as in Hibbins et al. (2023), and the Gaussian-Coalescent model with *λ* fixed at either 0 or 1. We did not attempt to estimate *λ* on this first analysis, as the datasets were small (3 values). Note that we used a corrected version of `seastaR`, as the one used in Hibbins et al. (2023) had an implementation error^1^ that led to unrealistically high estimates of the evolutionary rate (going as high as 2.7 *×* 10^5^, see Fig. 6 in Hibbins et al. 2023). As above, the variances were estimated using the REML procedure.

**Figure 7.**
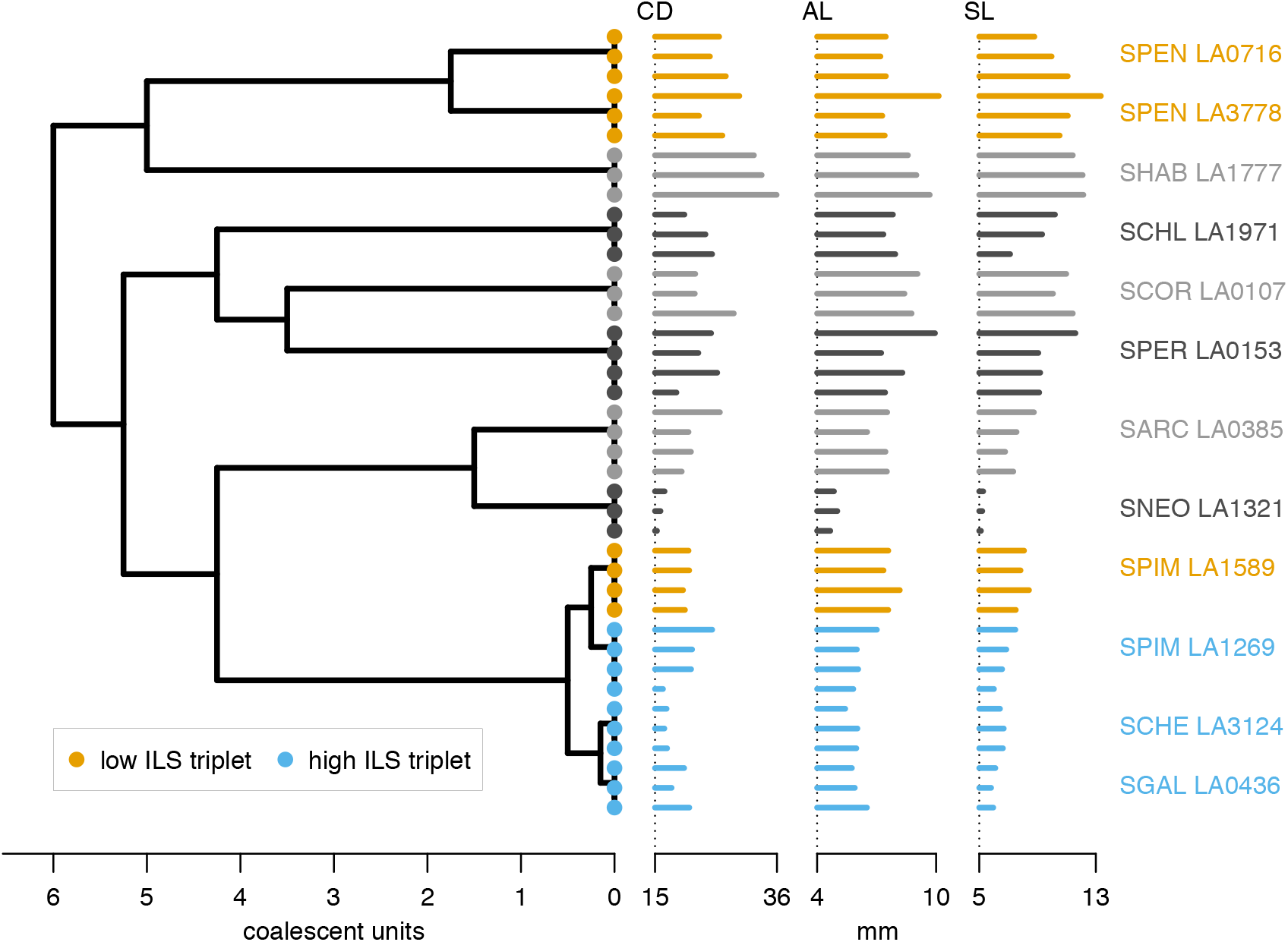
Phylogenetic tree of the wild tomato populations extracted from Pease et al. (2016) scaled in coalescent units as in Hibbins et al. (2023), with individual measurements (in mm) of corolla diameter (CD), anther length (AL) and stigma length (SL) from Haak et al. (2014). Species from “low ILS” (orange) and “high ILS” (blue) triplets are colored.

We then used the “full” dataset, that included as many populations and individual trait measurements as possible. We excluded two species from the tree that did not have any trait data (*S. huaylasense* and *S. tuberosum*), and removed three species IDs from the trait dataset that were not in the tree (SCER, SCHM and SLYC). Some species contained several populations (accessions). When this was the case, and when we did not have any further information on the divergence time between the different populations of a given species, we kept only the population with the highest number of samples. The resulting tree had 12 populations, with between 3 and 4 measurements per population, a total height of 6 coalescent unit, and room for relatively low overall ILS (see Fig. 7). We then compared the fit of seven models: the GC with *λ* either estimated, or fixed to a value of 0 or 1, either with or without additional within-population variation, and the BM with within-population variation, all permitted by the `R` implementation in `phylolm`. We compared the AIC weights (wAIC, Wagenmakers and Farrell, 2004) of each model and their estimated evolutionary rates (using REML). For each model, we also computed the total expected within-population variance. For the BM with extra within-population variation, this was simply the within-population variance parameter 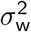. For the GC models, it was 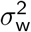 plus the evolutionary within-population heritable variance expected from the coalescent as computed in (20) with *t*_*u*_ = *T* the total height of the tree.

#### 3.3.2. Results

When studying the population mean traits on triplets, we found evolutionary variance rates qualitatively similar to those by Hibbins et al. (2023), with the BM model estimating 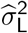 generally higher for the “high ILS” triplet compared to methods taking ILS into account, but similar for the “low ILS” triplet (Fig. 8). Further, in this setting where only triplets are used, our GC method with *λ* fixed to 1 (root at equilibrium) is equivalent to the `seastaR` model as explained in Remark 6, so that they give the exact same estimates 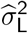. When fixing *λ* to zero (i.e., root variance v_0_ = 0), we found estimates that were in between, indicating that this model estimated a higher evolutionary variance compared to the model with *λ* = 1, but smaller compared to the BM, which accounted for no within-population variation at all.

**Figure 8.**
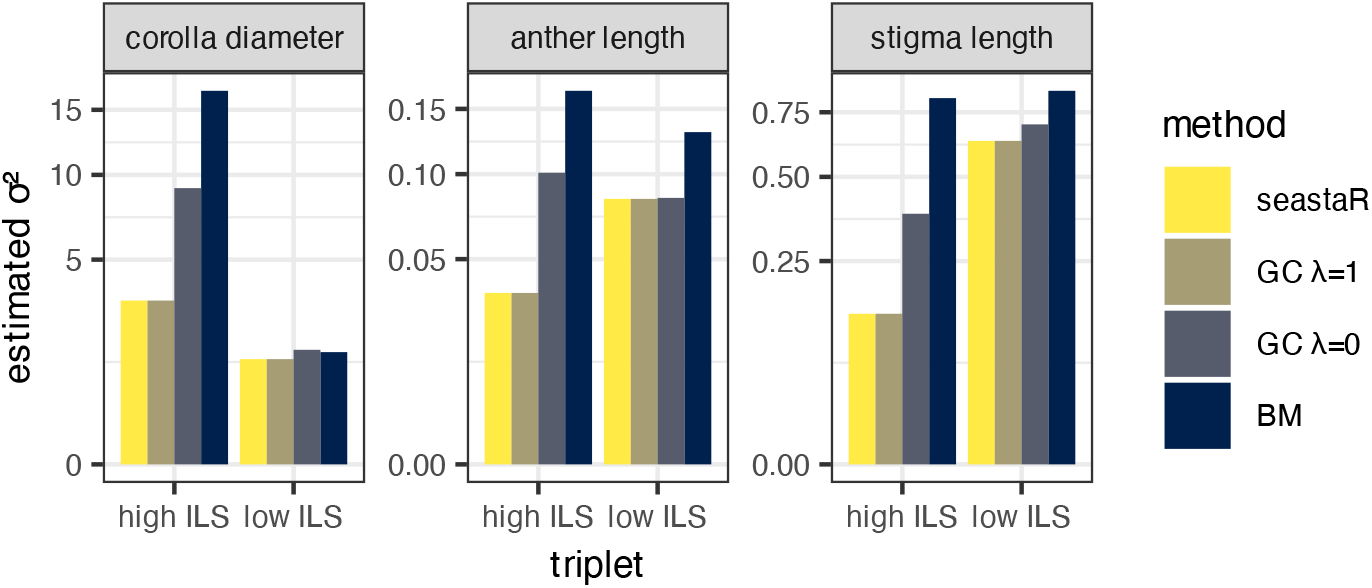
Estimated variance rate 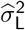 (displayed on the square-root scale) from population means, for three floral traits and three triplets as in Hibbins et al. (2023) (“low ILS”, *T* = 6 and “high ILS”, *T* = 0.5). Each panel uses a different limit for the vertical axis. Colors indicate the statistical method used: `seastaR` from Hibbins et al. (2023), the GC model with *λ* fixed at 0 or 1, and the BM. Under low ILS, 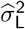 is similar across methods. Under high ILS, 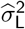 is generally higher for the BM model compared to ILS-based models.

When analyzing the full dataset with multiple values per population, we found that AIC favored the GC model with *λ* fixed and without extra non-evolutionary within-population variation (Fig. 9). The exact value of *λ* seemed to have less impact, with *λ* = 1 slightly favored for the corolla diameter and stigma length and *λ* = 0 for the anther length. This indicates that the GC model fits these data better than the BM model with added within-population variation. In other words, the evolutionary within-population variation expected from the coalescent was sufficient to explain the trait variability within each population.

**Figure 9.**
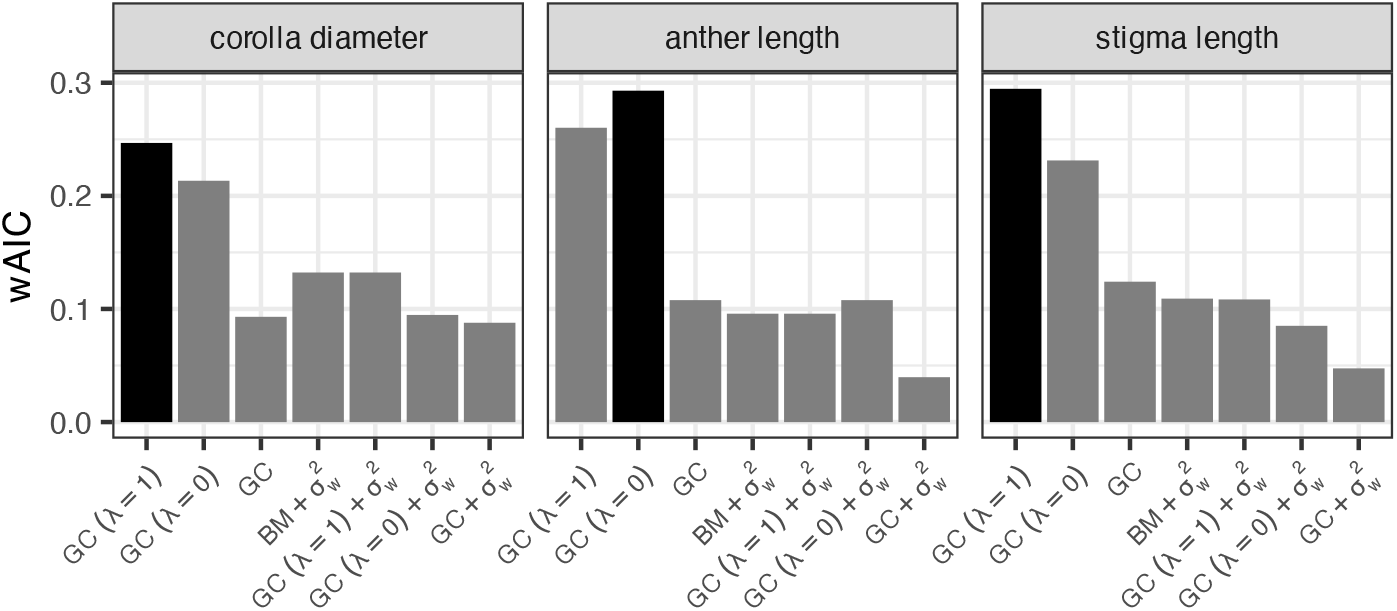
AIC weight for each model fitted to the full tomato tree, for the three floral traits, keeping replicate measurements within each populations. Models are the GC with *λ* fixed to 0 or 1 or estimated without or with an extra non-evolutionary withinpopulation variance parameter 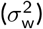, and the BM with within-population variance. The model in black is the one with the highest AIC value: it is always a GC model with fixed *λ* and no extra variation.

In most cases, the estimated evolutionary rate and the total within-population variation were highly similar across models, including the BM with extra within-population variation (Fig. 10). For corolla diameter, methods that included extra non-evolutionary variance tended to estimate a lower evolutionary variance.

**Figure 10.**
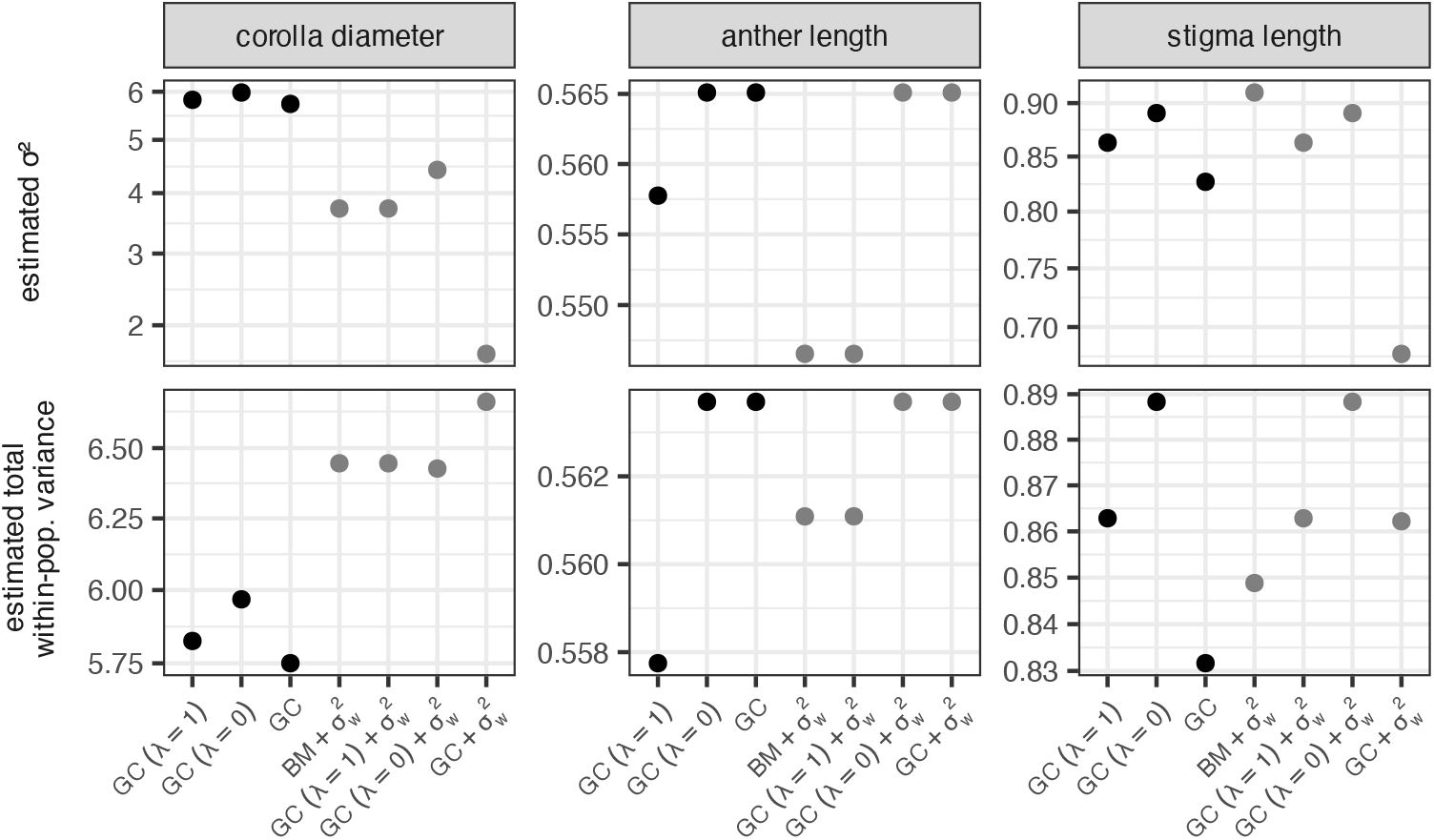
Estimated evolutionary variance 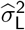 (top row) and total within-population variance for each model fitted to the full tomato tree (12 populations, 40 individuals). Variances are displayed on the square-root scale. Note that the scale for each panel is different, and often very narrow. Models are the GC with *λ* fixed to 0 or 1 or estimated and the BM, possibly with an extra non-evolutionary within-population variance parameter (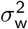, gray). Estimates are consistent between methods.

Note that, because the phylogeny is an ultrametric tree and we assumed a constant rate 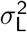, the within-population variance from the GC model, given in (20), is the same for all present-day populations. Further work could consider fitting a GC model with rates that differ in some clades or some branches, as permitted by Prop. 2, and as advocated recently in Mo and Hahn (2026).

## 4. Discussion

In this work, we describe a probability model for the evolution of a trait along a species phylogeny, in which the trait is controlled by one or several genes, that are themselves affected by incomplete lineage sorting, as modeled by the coalescent. If the phylogeny has reticulations, this process also includes the effect of gene flow, admixture, or hybridization. Our model accounts for ancestral polymorphism in the population at the root of the phylogeny, and implies an expected amount of heritable within-species variation in each present-day population. For tomato floral traits, this was sufficient to explain the within-population variation observed in the data.

Future work could survey other traits and groups to assess whether this pattern holds more generally.

Unlike prior methods accounting for ILS, our model is independent of taxon sampling, in the sense that the model for a sub-sample of populations remains unchanged when considering either the full or reduced phylogeny (given a fixed root population). Our model also applies to any phylogenetic network, of any topology and number of taxa, and applies to individual values for any number of individuals per taxon. It is implemented in two software packages (`phylolm` in `R` and `PhyloTraits` in `julia`) and can be fitted to data at no or little additional computational cost compared to standard PCM tools.

When the trait is polygenic controlled by many genes and if the effects of individual loci evolve with a finite variance rate, we showed that the distribution of the trait data is approximately Gaussian, leading to our Gaussian-Coalescent model.

In practice, there are several caveats for using our GC model. First, it requires a species phylogeny with branch lengths in coalescent units, which can be obtained as the ratio of number of generations over effective population size. Coalescent units can be estimated from the amount of gene tree discordance or from site pattern frequencies. While these estimates can be accurate at low values, such as between recently diverged populations, long coalescent units cannot be estimated accurately from genealogical discordance alone. This is because coalescence between individuals is extremely likely beyond 10 coalescent units, say, so that discordance disappears and becomes uninformative about the exact length of a branch in coalescent units, when it is long. For long coalescent times, accurate estimates may come from other units on branch lengths (substitutions, generations, calendar time) and assumptions on how these other units correlate with coalescent units. This issue has been considered for calibrating networks, although primarily to calibrate branch lengths proportional to time (Karimi et al., 2020; Tabatabaee et al., 2025; Xu and Ané, 2024).

Second, the GC model has one extra parameter compared to the Brownian motion: the trait variance in the root population. Ancestral polymorphism is typically not observed. Our simulations show that present-day data contain scant information about this ancestral variance, which is estimated with low precision. However, it is reasonable to assume that the root population was at equilibrium for the process (by fixing *λ* = 1). In simulations and in the tomato flower dataset, this choice was most often favored by AIC. It led to accurate estimates of the other parameters in most simulation scenarios. Under high ILS, it caused some bias in estimating the evolutionary rate, but much less bias and more accuracy than when using the BM model ignoring ILS.

Finally, our polygenic trait model assumes that the trait is the result of purely additive effects across genes, without epistatic interactions. Also, individuals are assumed haploids, carrying one copy of each locus. Our model readily applies to diploid individuals, if we further assume that the two copies of each locus have additive effects, like alleles across loci. Adding epistasis between loci and a model of dominance within loci would all be interesting future work. Valuable extensions also include relaxing the assumption that each locus evolves according to a Lévy process, modeling selection at the trait level such as with correlated loci (Bulmer, 1971), and considering the correlated evolution of several traits.

As described, our GC model should be used to analyze trait data at the individual level, because the GC variance includes within-population heritable variation. Averaging multiple values to estimate population means leads to a model violation, because taking the mean across individuals reduces the variance. This model violation also applies when using `seastaR`, which requires a single value per population. The tomato floral data analysis on triplets conducted here and in Hibbins et al. (2023) applies such trait averages within populations, so that its conclusions might be taken with caution. If measurement error, environmental variation or plasticity are expected to dominate genetic variance, then this practice could be beneficial despite its model violation. However, a sound alternative is to combine the GC model with extra non-heritable variation, as done here on the full tomato dataset.

Future work could build on our theoretical framework to explore using the GC model for a vast array of phylogenetic comparative methods, such as phylogenetic ANOVA and phylogenetic regression, or the detection of shifts in evolutionary rates. Many traits are inherantly multivariate (Clavel et al., 2015; Clavel and Morlon, 2020). Careful considerations of the biological mechanisms underlying the evolution of a multivariate polygenic trait would be required to extend the GC model to this setting. For instance, would all the traits be controlled by the same set of genes, or have only a subset in common? Building such a multivariate model could be an interesting direction for future work. Another valuable extension would consist in using observed gene trees, rather than gene trees expected from the coalescent, as pioneered by Hibbins et al. (2023), combined with conditioning on *ρ* as in the GC. A challenge is that gene trees are estimated with branch lengths in substitution per site, rather then calendar time or coalescent units. It may be necessary to account for substitution rate variation (if any) and to calibrate gene trees to embed them into the species phylogeny (tree or network) with comparable edge lengths, so as to identify the placement of the root population *ρ* in each gene tree. Recently, Mo and Hahn (2026) built on the *C*^∗^ framework using a sample of gene trees with edge lengths in coalescent units, that can be embedded into the species tree. They used this embedding to map phenotypic rates from the species tree onto gene trees. Modifying their approach by slicing gene trees at *ρ* would permit conditioning on the root population and obtain a method that is stable to taxon sampling.

## Funding

This work was supported in part by the “International Emerging Action” program of the CNRS, and by the National Science Foundation through grants DMS-2023239 (to C.A.) and DMS-1929284 while C.A. was in residence at the Institute for Computational and Experimental Research in Mathematics in Providence, RI, during the “Theory, Methods, and Applications of Quantitative Phylogenomics” program.

## Conflict of interest disclosure

The authors declare that they comply with the PCI rule of having no financial conflicts of interest in relation to the content of the article. The authors declare the following non-financial conflict of interest: C.A. is a recommender for PCI Evolutionary Biology and for PCI Mathematical and Computational Biology.

## Scripts and code availability

The GC model is available as part of the `R` package `phylolm` (Ho and Ané, 2014), version 2.7.0; and of the `julia` package `PhyloTraits` (Bastide et al., 2018), version 1.2.0. The code to reproduce the simulations, and the data and code to reproduce the analysis of the tomato floral traits are available at https://github.com/cecileane/GCmodel-2026-data-code and at Zenodo (Bastide and Ané, 2026).

## A. Graphical model and proof of covariance formulas

### A.1. Graphical model

In this section we express the one-locus model in Def. 3 as a formal graphical model. To this end, for each node in the network we consider two individuals drawn at random from the population that this node represents.

#### Definition S1

(Graphical model). Let *N* be phylogenetic network. For each node *u* ∈ *V* (*N*), we define two random variables *Y*_*u*,1_ and *Y*_*u*,2_. The joint distribution of the vector **Y** = (*Y*_*u,i*_)_*u*∈*V* (*N*),*i*∈{1,2}_ is defined by the following graphical model. At the root *ρ* of *N*, we assume that *Y*_*ρ*,1_ and *Y*_*ρ*,2_ are independent and identically distributed according to a prior distribution *P*_0_.

Let *u* be a non-root node of, with *m* ≥ 1 parents *p*_1_, · · ·, *p*_*m*_, edge (*p*_*k*_, *u*) of length *ℓ*_*k*_ in coalescent units, inheritance probability *γ*_*k*_, and centered Lévy process 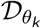. Further, let *H*_1_ and *H*_2_ be two independent random variables such that ℙ [*H*_1_ = *k*] = ℙ [*H*_2_ = *k*] = *γ*_*k*_ for all 1 ≤ *k* ≤ *m*. Define a random variable *T* as follows. If *H*_1_ ≠ *H*_2_, then *T* = + ∞. If *H*_1_ = *H*_2_ = *k*, then *T* is drawn from an exponential distribution with mean 1, that is, ℙ [*T ≤ ℓ*|*H*_1_ = *H*_2_ = *k*] = *q*(*ℓ*). Finally, define a random variable *J* such that ℙ [*J* = 1] = ℙ [*J* = 2] = 1*/*2. The distribution of (*Y*_*u*,1_, *Y*_*u*,2_), conditional on all parent variables 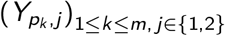 and on *H*_1_, *H*_2_, *T* and *J* is then defined as follows.

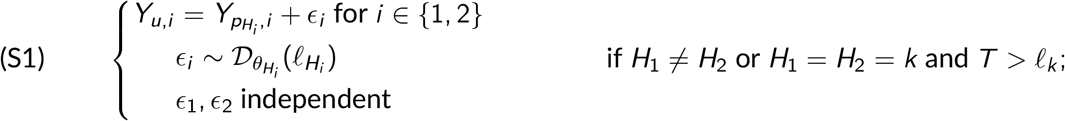

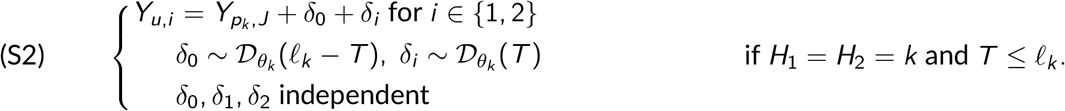

For convenience, we denote by *C*_*u,k*_ the random event *H*_1_ = *H*_2_ = *k* and *T* ≤ *ℓ*_*k*_, and by 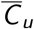 the complement over all *k*: *H*_1_ ≠ *H*_2_ or *H*_1_ = *H*_2_ and 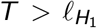. Note that, using the properties of the Lévy process, 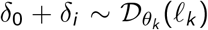 in (S2) has the same distribution as *ϵ*_*i*_ in (S1) and does not depend on *T* .

**Proposition S1**. *For any node u in N, the random variables Y*_*u*,1_ *and Y*_*u*,2_ *in Def. S1 are exchangeable. Further, consider a non-root node u, any i, j* ∈ *{*1, 2*}, and use notations in Def. S1. Conditional on H*_1_, *H*_2_, *T, J and on H*_*i*_ = *k, Y*_*u,i*_ *is distributed as* 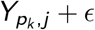 *where* 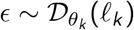 *is independent of* 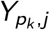.

*Proof*. We prove the exchangeability by induction on nodes in *N*, considering them in preorder (from root to leaves). By definition, *Y*_*ρ*,1_ and *Y*_*ρ*,2_ are independent and identically distributed, and hence are exchangeable. Let *u* be a node in *N*. The exchangeability of *Y*_*u*,1_ and *Y*_*u*,2_ derives from the symmetry in *i* ∈ {1, 2} in (S1) and (S2), using the exchangeability of (*H*_1_, *H*_2_), the induction hypothesis on 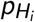 for (S1), and the symmetry of ℙ [*J* = 1] = ℙ [*J* = 2] for (S2).

We now turn to the second claim and fix *i, j* ∈ *{*1, 2*}*. Conditional on *H*_*i*_ = *k* and on 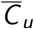, 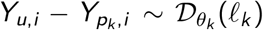 from (S1), regardless of *T, J* and *H*_*i*_*′* for *i*^*′*^ ≠ *i*. The distribution of *Y*_*u,I*_ conditional on *H*_*i*_ = *k* and 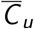 is then identical to that of 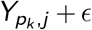 by exchangeability of 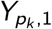 and 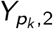. Conditional on *C*_*u,k*_ and on 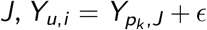 with 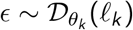 from (S2) and properties of the Lévy process. Since 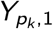 and 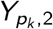 are exchangeable, the distribution of *Y*_*u,i*_ conditional on *C*_*u,k*_ and *J* is the same as that of 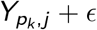, regardless of the value of *J* . Note that as a corollary, the claim holds conditional on *H*_*i*_ = *k* only (e.g., integrating over *T* and *J*). □

#### Proposition S2

*Assume that Y evolves on N according to the model in Def. 3 for one locus. Consider two individuals sampled from each population u (ancestral or extant) in N, with values Y*_*u*,1_ *and Y*_*u*,2_. *Then the joint distribution of* (*Y*_*u,i*_)_*u*∈*V* (*N*),*i* ∈ *{*1,2}_ *is equal to the joint distribution given by the graphical model in Def. S1*.

*Proof*. We use induction on the nodes in *N*, listed in a preorder, that is: the root comes first and any node *u* must be listed after all its parents have already been listed. We note *u < v* if *u* is listed before *v* . We will prove by induction on *v*, that the joint distribution of (*Y*_*u,i*_)_*u*≤ *v,i* ∈ {1,2}_ (up to *v*), is equal under Def. 3 and Def. S1. By definition, *Y*_*ρ*,1_ and *Y*_*ρ*,2_ are independent and identically distributed from *P*_0_ in both models.

Now assume that the joint distribution up to *v* are identical, and take *u* the following node in the preorder. Take the two individuals 1 and 2 in population *u*, and define *G*_*p*_ as the tree for these two individuals, for the locus of interest, truncated to the parts that evolved within the *m* parent populations (*p*_*k*_, *u*) for *k* = 1, …, *m*. This gene tree includes the identity *H*_*i*_ of the parent population that individual *i* ∈ {1, 2} was inherited from (at the locus being considered). For example, if individual 1 came from *p*_1_, then *H*_1_ = 1. By definition of the network multispecies coalescent process (Degnan, 2018; Fogg et al., 2023), ℙ [*H*_*i*_ = *k*] = *γ*_*k*_, and these variables coincide with the *H*_*i*_ introduced in Def. S1. Individuals 1 and 2 can either coalesce before reaching a parent of *u*, or not coalesce, in which case *G*_*p*_ consists of a “forest” of two separate gene edges. We denote by *T* the coalescent time (starting from *u*) between individuals 1 and 2 at that locus. *T* has an exponential distribution, and coincide with the variable introduced in Def. S1.

Individuals 1 and 2 do not coalesce before reaching a parent of *u* if either *H*_1_ ≠ *H*_2_ (they came from different parents) or if *H*_1_ = *H*_2_ = *k* and *T > ℓ*_*k*_, that is, if event 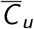 from Def. S1 occurs. In this case, each individual 1 and 2 chooses its own ancestor in its own parent population 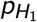 and 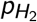, with traits 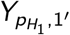 and 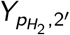, where *i*^*′*^ the ancestor individual of *i* . From these ancestors, they evolve as the Lévy process, so that 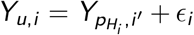, with 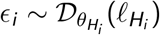 independent from the rest of the variables. Individual *i*^*′*^ might not be sampled at node 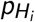, as we assumed that only two individuals were sampled per population. However, since the backward-in-time coalescent process samples the ancestor of *i*^*′*^ at random from the parent population 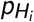, this ancestor’s trait has the same distribution as 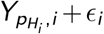, with *i* ∈ *{*1, 2*}*. We then recover exactly (S1) from Def. S1.

Finally, individuals 1 and 2 coalesce if *H*_1_ = *H*_2_ = *k* and *T* ≤ *ℓ*_*k*_, i.e., event *C*_*u,k*_ from Def. S1 is true. In this case they both have the same ancestor, picked at random in parent population *p*_*k*_ . Without loss of generality, we can take this ancestor to be *J*, with ℙ [*J* = 1] = ℙ [*J* = 2] = 1*/*2. Then the two individuals diverge after a time *ℓ*_*k*_ − *T* from *p*_*k*_, and, according to the Lévy model of evolution, we get, for 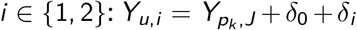, with 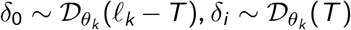, and *δ*_0_, *δ*_1_ and *δ*_2_ mutually independent and independent from other variables. We hence recover (S2) from Def. S1.

To summarize, the joint distribution of all trait variables are equal at all nodes up to *u* in preorder, which ends the proof. □

*Remark* S1 (Gaussian marginals). In the special case when 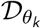 is a centered Gaussian distribution, the graphical model is not jointly Gaussian, due to the random variables *H*_*i*_ and *T* that are not Gaussian. Marginally, when considering the locus effect of a single randomly sampled individual, *Y*_*u,i*_ is not generally Gaussian either. See appendix B for examples. However, as *ϵ*_0_ + *ϵ*_*i*_ ∼ Normal 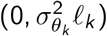 does not depend on *T*, then *Y*_*u,i*_ is Gaussian conditional on its ancestor *Y*_*v,J*_ if *u* is a tree node with unique parent *v* . Consequently, *Y*_*u,i*_ is Gaussian conditional on the locus gene tree history, with variance 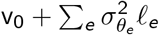 where the sum is over edges *e* that the gene tree evolved in. For example, if *N* is a tree, then each *Y*_*u,i*_ is marginally Gaussian, even though the full vector of all variables is not a Gaussian random vector. This property is maintained if the network is time-consistent in variance units *σ*^2^*ℓ*, that is, if all paths from the root to *u* have equal lengths.

#### Remark S2

(Mutational model). When 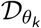 is defined by a compound Poisson process, the model is similar to that described in Schraiber and Landis (2015) (who focused on a single evolving population), with the same propagation rules. Here we model ancestral polymorphism with distribution *P*_0_ in the root population *ρ*, whereas Schraiber and Landis (2015) assume that the trait of all individuals in *ρ* is fixed at zero.

### A.2. Proofs of main results

In this section, we assume that the increment distribution has a finite variance rate, and we present the proofs delayed to this appendix. As before, we omit *ρ* to simplify notations, but all expectations, variances and covariances are calculated conditional on the trait distribution in *ρ*.

*Proof of Proposition 2 (Recursion formula)*. Thanks to Prop. S2, the quantities in Prop. 2 are the mean *E*_*u*_ = E [*Y*_*u,i*_], covariances Ω_*u,v*_ = ℂov [*Y*_*u*,1_; *Y*_*v*,2_] and variances Φ_*u*_ = Var [*Y*_*u*,1_], all conditional on the distribution *P*_0_ in the root population *ρ*, where **Y** = (*Y*_*u,i*_)_*u*∈*V*(*N*),*i* ∈ {1,2}_ are the random variables defined by the graphical model in Def. S1.

As (4) is a special case for a tree node with *m* = 1 parents, we only need to prove (5). Consider node *u* and assume that (3)-(5) hold for its parent nodes. Using notations from Def. S1 we get by Prop. S1 that

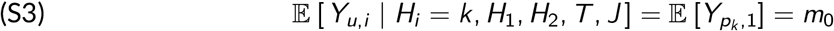

where the second equality comes from (3) at parent node *p*_*k*_ . This is a stronger version of (3) at *u*, since it implies that E [*Y*_*u,i*_] = E [*m*_0_] = *m*_0_, that is, (3) at *u*. For variances and covariances, we condition again on the partial gene tree history *G*_*u*_ = {*H*_1_, *H*_2_, *T, J}* describing how *u* inherits from its parents. First,

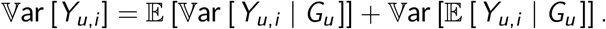

By (S3), the second term is 0. For the first term, using Prop. S1, ℙ [*H*_*i*_ = *k*] = *γ*_*k*_ and the variance of the Lévy process, we get:

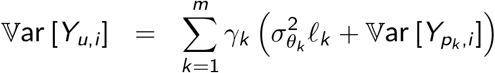

as claimed in (5a). Next, for individuals 1 and 2 in *u*, we condition again on the partial gene tree history *G*_*u*_ = *{H*_1_, *H*_2_, *T, J}* at *u*:

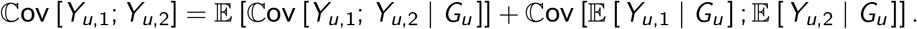

By (S3) again, the second term is 0. For the first term, we condition more specifically on *C*_*u,k*_ for *k* = 1, …, *m* or their complement 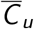. If 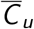 occurs, then from (S1) the increments 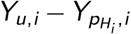 are independent (*i* = 1, 2) and their correlation is 0 conditional on *G*_*u*_ and 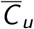. If instead *C*_*u,k*_ occurs, then from (S2) the increments are 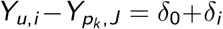 so their covariance is 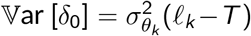. Therefore, we get

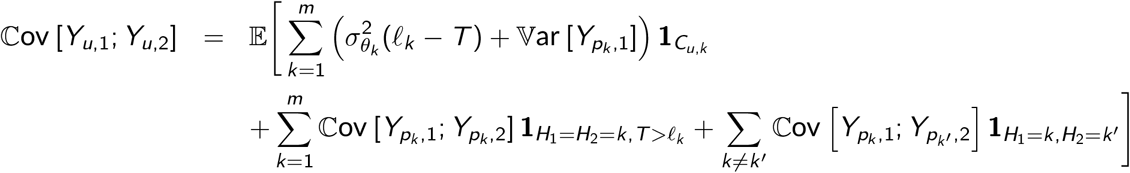

from which (5b) follows. Finally, if *v* is not a descendant of *u*, then the increment between *Y*_*u,I*_ and its parent is independent of *Y*_*v,j*_ . Conditioning on *G*_*u*_ and using (S3) again, we get

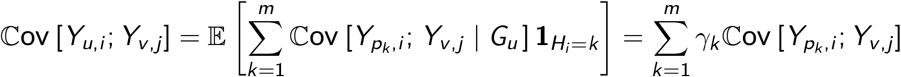

which gives (5c) and finishes the proof. □

Next, we turn to proving Prop. 3.

*Proof of Proposition 3 (Sampling stability)*. If networks are allowed to have nodes of degree 2 and if we do not seek to suppress any degree-2 node in *N*_−*t*_ when pruning taxon *t* from *N*, then Prop. 3 follows directly from Prop. 2, as the recursion formulas for a given node only depends on its parent nodes, and not its sibling nodes.

Now assume a constant variance rate 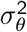 on all the branches. We will show that this assumption allows for the suppression of a node *v* along a lineage *u* → *v* → *w*, if *v* is of degree-2 (*u* and *w* are its only neighbors), by fusing (*u, v*) and (*v, w*) into a single edge of length *ℓ*_*uw*_ = *ℓ*_*uv*_ + *ℓ*_*vw*_ coalescent units, and inheritance *γ*_*uw*_ = *γ*_*vw*_ .

Take any leaf taxon *t* in the network. Being a leaf, it has a unique parent *v* . First assume that *v* is a tree node. If *v* is the root *ρ*, then we just keep this node, so that the the conditioning used for all expectations is not affected. Otherwise, *v* has a unique parent *u*. If *v* has more than two children in *N*, then *v* is still a tree node with at least 2 children in *N*_−*t*_, and the formulas of Prop. 2 are unchanged in *N*_−*t*_. If *v* has exactly two children in *N, t* and *w*, then *v* has degree two after removing *t*, with parent *u* and child *w* . Node *w* can be a leaf or internal, possibly hybrid. Assume that *w* has *m* + 1 parents in *N*, that are *v* and (*p*_*k*_)_1≤*k* ≤*m*_ (if *w* is a tree node then *m* = 0). Then from (5a), (5b) and (5c) in the full network:

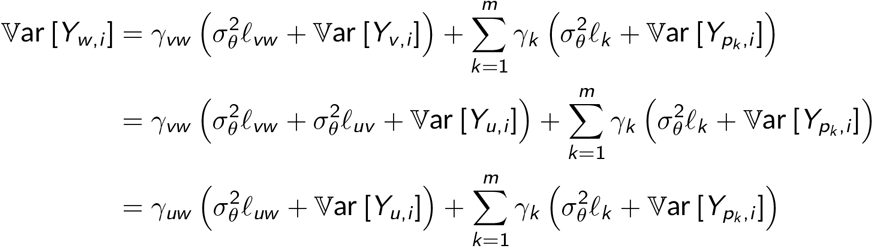

where *γ*_*uw*_ = *γ*_*vw*_ and *ℓ*_*uw*_ = *ℓ*_*uv*_ + *ℓ*_*vw*_ are for the fused edge (*u, w*) after suppressing *v* . Similarly:

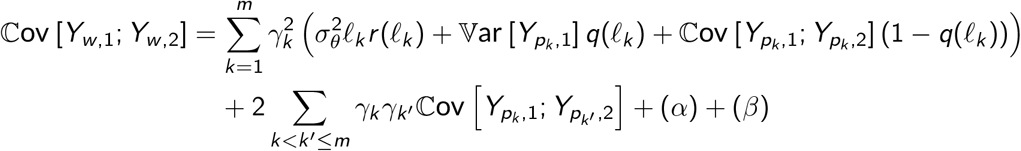

where

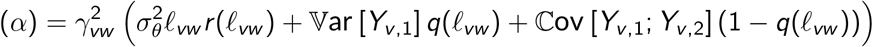

and

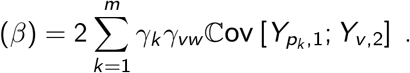

As *v* is a tree node, 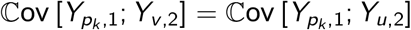 for any *k*, and *v* can be replaced by *u* in (*β*). Also, still using that *v* is a tree node:

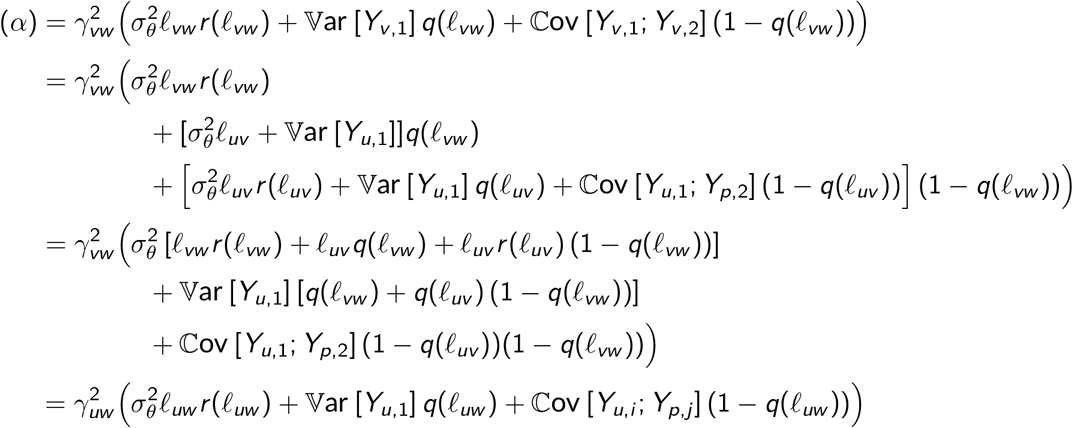

and we recover formulas (5a), (5b) and (5c) applied to the branch going directly from *u* to *w*, skipping node *v* of degree two, that can be suppressed, yielding subnetwork *N*_−*t*_. This ends the proof in the case when *v* is a tree node.

Now assume that *v* is a hybrid node. If it has two or more children, then it is still a hybrid node with at least 1 child in *N*_−*t*_, and the formulas of Prop. 2 are again unchanged in *N*_−*t*_. Otherwise *t* is the only child of *v* in *N*, and when *t* is pruned, *v* is removed along with the edges from *v* ‘s parents *p*_*k*_ to *v* . Then, each *p*_*k*_ in the subnetwork is either of degree more than two, in which cases it remains, or exactly two, in which case it can be suppressed using the same derivations as above, or exactly one, i.e. it is the root *ρ*, that we preserve in the subnetwork. This ends the proof, as suppressing all nodes of degree exactly two in the subnetwork leads to the same propagation formulas on the subnetwork. □

*Proof of Proposition 5 (BM on a rescaled network)*. We re-write covariances in Cor. 4 so that they have the form in (9). To prove (10), we write:

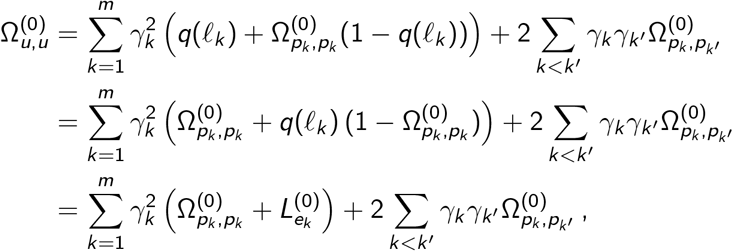

which takes form (9) using the rescaled lengths *L*^(0)^. The proof of (11) is similar.

Next, take nodes *u* → *v* → *w* along one lineage, such that *v* is a tree node and *ℓ*_*uv*_ +*ℓ*_*vw*_ = *ℓ*_*uw*_ .

Then:

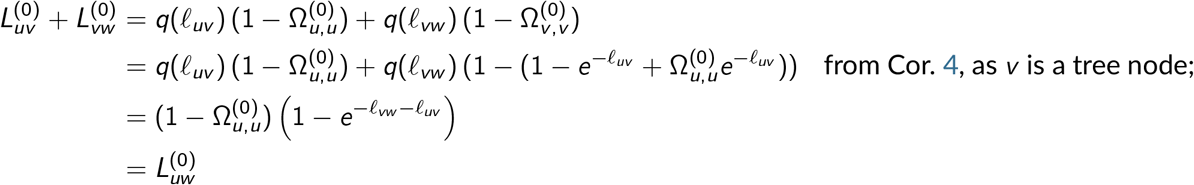

so that the branch length transformation (10) is additive. Similarly:

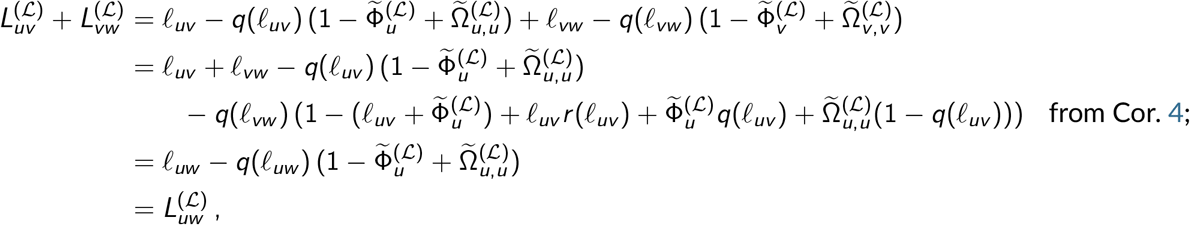

where we used the relationship: 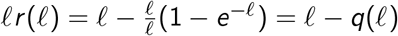. □

The following extends (11) in Prop. 5 to the case when the rate is not constant. It shows that **Ω**^(ℒ)^ can still be seen as a BM on the phylogeny, although the transformed branch lengths now depend on the branch parameters 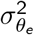.

#### Proposition S3

(BM on a rescaled network variable rate). *The variance matrix* **Ω**^(ℒ)^ *is the same as the one obtained from a BM on the phylogeny, with branch-specific variance parameters* 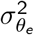 *and rescaled branch lengths:*

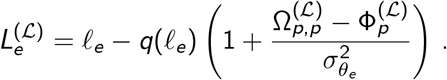

*Proof*. As in the proof of Prop. 5, we simply re-arrange terms in (7) from Cor. 4 to recover form (9). □

*Proof of Proposition 6 (Closed-form expression in the tree case)*. We use the recursions from Cor. 4 to prove (12) and (14) by induction on nodes *u, v* listed in preorder. At the root, *t*_*ρρ*_ = 0, so that 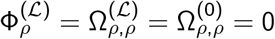 and (12) and (14) hold.

Now assume that (12) and (14) hold for all nodes listed before *u* in preorder, and for its parent node *p* in particular. To prove (14a), we use (7a), that gives 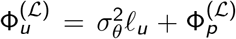 where *ℓ*_*u*_ is the length of the branch from *p* to *u* in coalescent units. By induction, 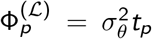 so we get 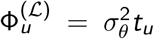 from *t*_*u*_ = *t*_*p*_ + *ℓ*_*u*_. To prove (12) and (14b), first consider a population *v*≠ *u* listed before *u*. From (7c) and (8a), we get 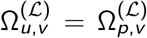 and 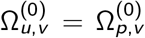. As *v* is not a descendant of *u* by assumption, *t*_*uv*_ = *t*_*pv*_, and we get (12) and (14b) by induction. Now consider the case *u* = *v* . Applying (8a) and by induction, we get 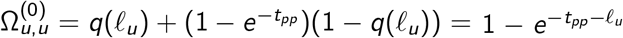, which is precisely (12) as *t*_*uu*_ = *t*_*pp*_ + *ℓ*_*u*_. Finally, applying (7b) and by induction, we get 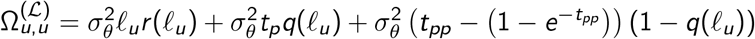. Using the relationships 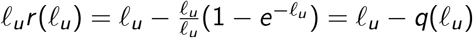, and again *t*_*uu*_ = *t*_*pp*_ + *ℓ*_*u*_, we get (14b) as expected.

The branch length transformations (13) and (15) then directly follow from Prop. 5. □

### A.3. Mean population trait evolution on a network

Recall that *M*_*u*_ = E_*i*_ [*Y*_*u,i*_] denotes the mean trait among all individuals *i* in population *u* for a given locus, and *V*_*u*_ = Var_*i*_ [*Y*_*u,i*_] = E_*i*_ (*Y*_*u,i*_ − *M*_*u*_)^2^ its variance within population *u*. Then the moments of *Y*_*u,i*_ are linked to those of *M*_*u*_ and *V*_*u*_ as claimed in Lemma 7, which we prove below.

*Proof of Lemma 7*. We obtain (16) by re-writing the variances and covariances in a different order. In the following, we use E_coal_ to specify that we take the expectation over the coalescent process, and E_*i*_ for the expectation over individuals within a population. With no index, the expectation is taken over the whole process. First, it is easy to see that

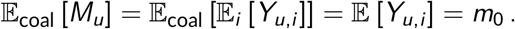

For the covariance, we get:

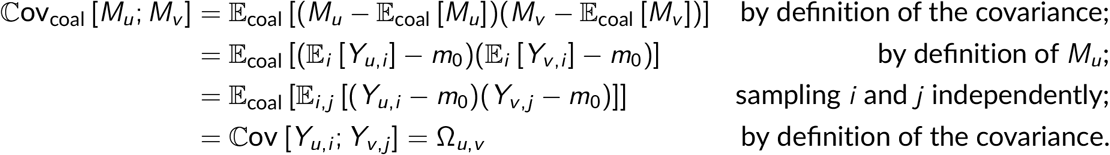

Using similar techniques, we get for the variance:

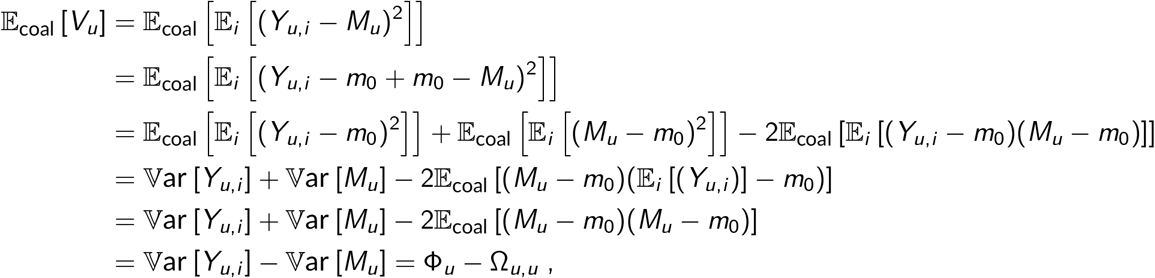

which ends the proof. □

## B. Non-Gaussian joint distribution

We give here a numerical illustration of Remark S1, which notes that the trait distribution may not be Gaussian given a finite number of loci, even if increments have a Gaussian distribution.

We simulated a trait under the network in Fig. S1, controlled by *L* = 1 locus whose effect evolved as a Brownian motion with rate 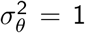, starting with variance v_0_ = 0.1 at the root *ρ* (*λ* = 0.1). Some of these settings are fairly extreme to obtain non-Gaussian distributions, to be ‘far’ from the the Gaussian limit (Prop. 10). Namely, we chose a small *L* = 1, and a phylogeny that is not time-consistent (in variance units) to obtain non-Gaussian marginals. For example, the two paths from the root to the hybrid node have very different lengths: 1.4 and 0.1.

**Figure S1.**
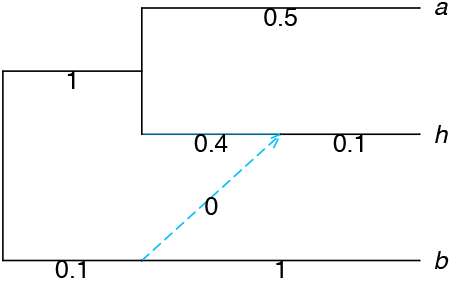
Network with branch lengths showing coalescent units, used to simulate the traits shown in Fig. S2. Both hybrid edges were assigned inheritance *γ* = 0.5. In each population, 2 individuals were simulated.

We simulated the trait for 2 individuals per population 100,000 times (Fig. S2). As population *a* is not below any reticulation, each individual *a*_1_ and *a*_2_ has a trait that is marginally Gaussian (histograms not shown). But their joint distribution is clearly not bivariate Gaussian, with non-elliptical contours (Fig. S2 left). For individuals *h*_1_, *h*_2_ in population *h*, their joint distribution is even more clearly non-Gaussian (Fig. S2, middle). This is due to the additional variation in gene trees inherited from the different parents at the reticulation. For individual *h*_1_ (or *h*_2_) taken individually, its trait is not Gaussian (Fig. S2, right), although mildly so.

**Figure S2.**
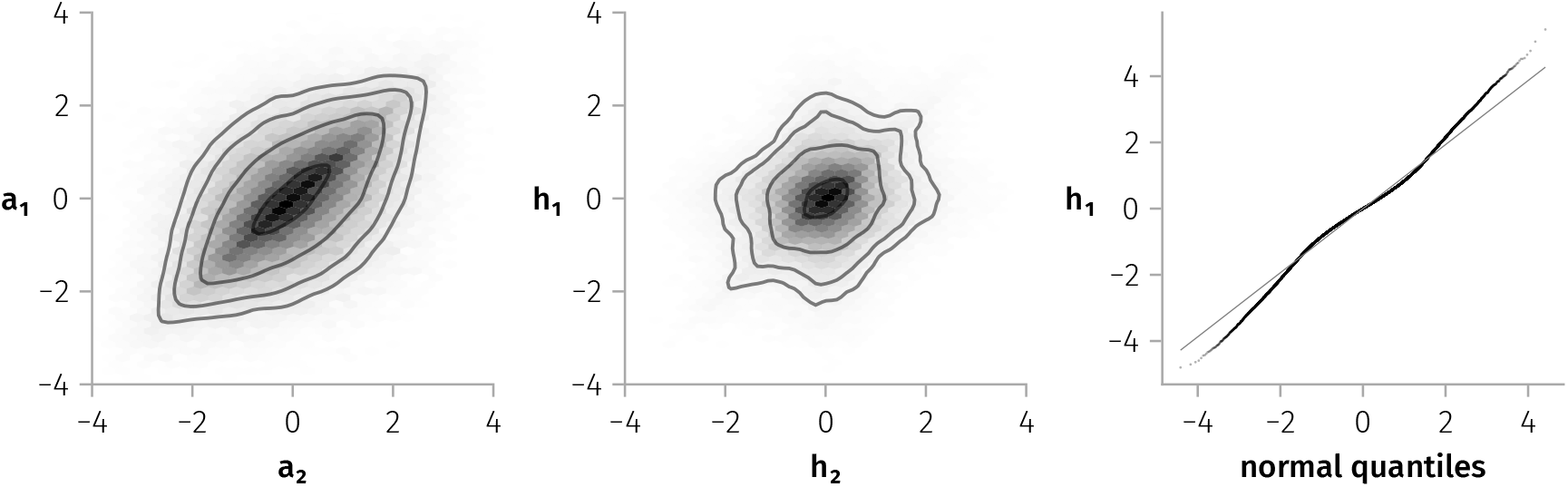
Joint trait distribution from *L* = 1 locus for two individuals *a*_1_ and *a*_2_ from population *a* (left panel), or *h*_1_ and *h*_2_ from population *h* (middle panel), under the network in Fig. S1. For each of *a*_1_ and *a*_2_, the trait followed a Gaussian distribution marginally (not shown). For *h*_1_ (and *h*_2_, by symmetry), the trait did not follow a Gaussian distribution marginally, as its quantiles do not align with those of a normal distribution (right panel).

## C. Limits in small or large populations

We assume here that *N* is a tree, and study the limits of the GC and *C*^∗^ covariances when the haploid effective population size 2*N*, assumed constant over *N*, becomes very small or very large. These two limits correspond to “no ILS” with immediate coalescence when *N* → 0, and “maximum ILS” with no coalescence until the root *ρ* when *N*→ ∞ .

For the GC covariance, we can re-write (18) and (20) from the main text with time in generations instead of coalescent units. We get

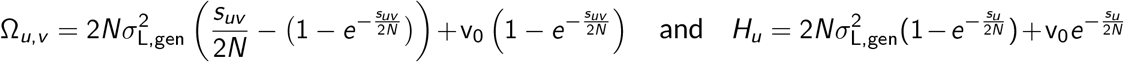

where 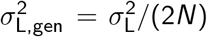 is the evolution rate per generation, *s*_*uv*_ = 2*Nt*_*uv*_ is the shared evolutionary time between *u* and *v* in generations, and *s*_*u*_ = *s*_*uu*_ = 2*Nt*_*u*_ is the number of generations from the root to *u*. It is then possible to study the limits of these equations when the population becomes very large or very small, either fixing 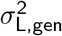 or 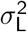:

- Under no ILS (*N* → 0), the model is equivalent to a BM model on the original tree as

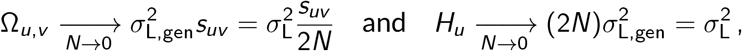

with an added within population variation of 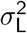 when the rate in coalescent unit is fixed rather than the rate in generations. Note that the variance within the root population v_0_ is “forgotten” at the limit.
- Under maximum ILS (*N*→ ∞), the model is equivalent to a single population (or star tree) with independent individuals, as

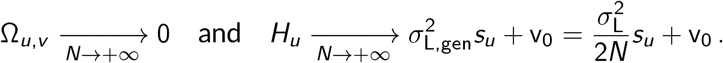

Note that, when varying *N*, the total variance of an individual sampled from population *u* remains unchanged, equal to 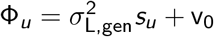, with time *s*_*u*_ in generations.

For the *C*^∗^ matrix, we first write (22) from the main text with time in generations:

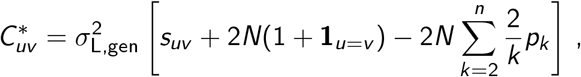

and change *N* to vary the level of ILS in the method by Mendes et al. (2018).

- →Under no ILS (*N* → 0), *C*^∗^ converges to the simple BM covariance matrix.
- Under maximum ILS (*N* → ∞), the covariances and variances in *C*^∗^ increase indefinitely. This is because the lineages never coalesce, and the process is conditioned on the individual’s locus effect at the last coalescent event, which is infinitely far in the past.

Note that Mendes et al. (2018) characterize “no ILS” and “maximum ILS” differently than by varying *N* (see Section 2.3.1 of their Appendix). For “no ILS”, they set *p*_2_ = 1 and *p*_*k*_ = 0 for *k >* 2, which corresponds to no ILS after the root *ρ* in *N* and a regular coalescent process before *ρ*, in which case their formulas converge to the simple BM with an extra term of 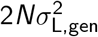 for the variance of the tip traits. For “maximum ILS”, they set *p*_*n*_ = 1 and *p*_*k*_ = 0 for *k < n*, which corresponds to no coalescent event after the root and a regular coalescent process before *ρ*, in which case their formulas converge to a simple BM with an extra variance 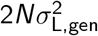 at the tips, and an extra covariance term 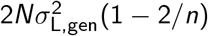 (1 2 *n*) for all pairs of tips.

## D. Supplementary figures

### D.1. Numerical experiments with seastaR

#### D.1.1. Methods

As detailed in Section 2.6, the package `seastaR` (version 0.1.0 at commit b23476f) is based on Hibbins et al. (2023) and Mendes et al. (2018), and contains two main functions:

- `get_full_matrix` takes a species tree as input, and implements the formulas of Mendes et al. (2018) on triplets. For two tips in the tree, their covariance is computed as the average covariance of these two tips across all the triplets containing them.
- `trees_to_vcv` takes as an input a set of gene trees relating the populations under study, with branch lengths assumed to be on the same scale (such as coalescent units). It calculates a covariance matrix from each gene tree and returns their average.

In Section 2.6.2, we focused on `get_full_matrix` because it takes the same input as the GC model: the species phylogeny. However, `get_full_matrix` is based on theory for 3-taxon trees, it has not been thouroughly evaluated for trees with more than three species; and it currently uses all triplets from the tree including those that do not span the root. An alternative approach recently described in Mo and Hahn (2026) can apply to a species phylogeny with more than three taxa:

1. simulate *N*_*g*_ gene trees from the (network) multispecies coalescent within the input species phylogeny; and
2. use these simulated gene trees as input to `trees_to_vcv`.

Currently, this approach is not natively supported by `seastaR`. It requires some light engineering to call an external program to simulate gene trees in step (1).

We reproduced the numerical experiments of Section 2.6.2 using this procedure, simulating *N*_*g*_ = 1 000 gene trees for each species tree used the `julia` package `PhyloCoalSimulations.jl` (v.1.1.0, Ané et al., 2025; Fogg et al., 2023). We also tracked the computation time of each covariance calculation method, after gene trees were simulated (that is, excluding the time to run step (1) above). All scripts are available on the companion repository https://github.com/cecileane/GCmodel-2026-data-code.

#### D.1.2. Results

Figures S3 and S4 reproduce the analyses presented in the main text on Figures 3 and 4, with added results using the `trees_to_vcv` procedure (colored full lines). First, and as expected from Mendes et al. (2018), both `seastaR` procedures give the same results on 3-taxon trees (Figure S3, dark purple). The difference between `get_full_matrix` and `trees_to_vcv` appears for larger trees, and deepens when the number of taxa increases.

Second, the linear decrease with *t* for the variance of A or the covariance between A and B (Fig. S3 and S4, left-most panel) mostly disappears with the `trees_to_vcv` method. This pathological behavior aligns with the problem mentioned by M. Hahn and M. Hibbins in their reviews (see Rogers, 2026) that `get_full_matrix` should only average over triplets that span the root. If a future version of `seastaR` implemented this modification, this would not change the behavior of the method for the variance of C or the covariance of A and C (Fig. S3, middle and right panels), as all the triplets containing C and two other tips span the root in the left tree of Fig. 2. The same goes for the covariance of C and D in the right panel of Fig. S4. Yet, `get_full_matrix` and `trees_to_vcv` still return different *C*^∗^ matrices in this case, so further improvements are needed for the two methods to return matching results.

The variances and covariances given by `trees_to_vcv` still depend on both the number of “extra” taxa and on the relative internal height *t*. This dependence is even stronger than that of `get_full_matrix` for (co)variances involving taxon C (for which all triplets span the root, see above). This is particularly visible, for instance, for the covariance of A and C (Fig. S3, right panel), that sees a sharp increase when *t* goes to 0. This could be a consequence of the conditioning of `trees_to_vcv` on the random coalescent time of each gene tree, as a smaller internal branch length leads to an average longer time of shared evolution between A and C. In contrast, the covariance between A and C remains constant and equal to zero for the GC method that condition on the species tree root population, and does not average over covariances computed using different conditioning times. A similar pattern can be observed for the covariance of C and D (Fig. S4, right panel), where `get_full_matrix` gives a constant covariance that is equal to the covariance computed by the GC method, but the covariance computed with `trees_to_vcv` exhibits some variations both in *n* and *t*.

**Figure S3.**
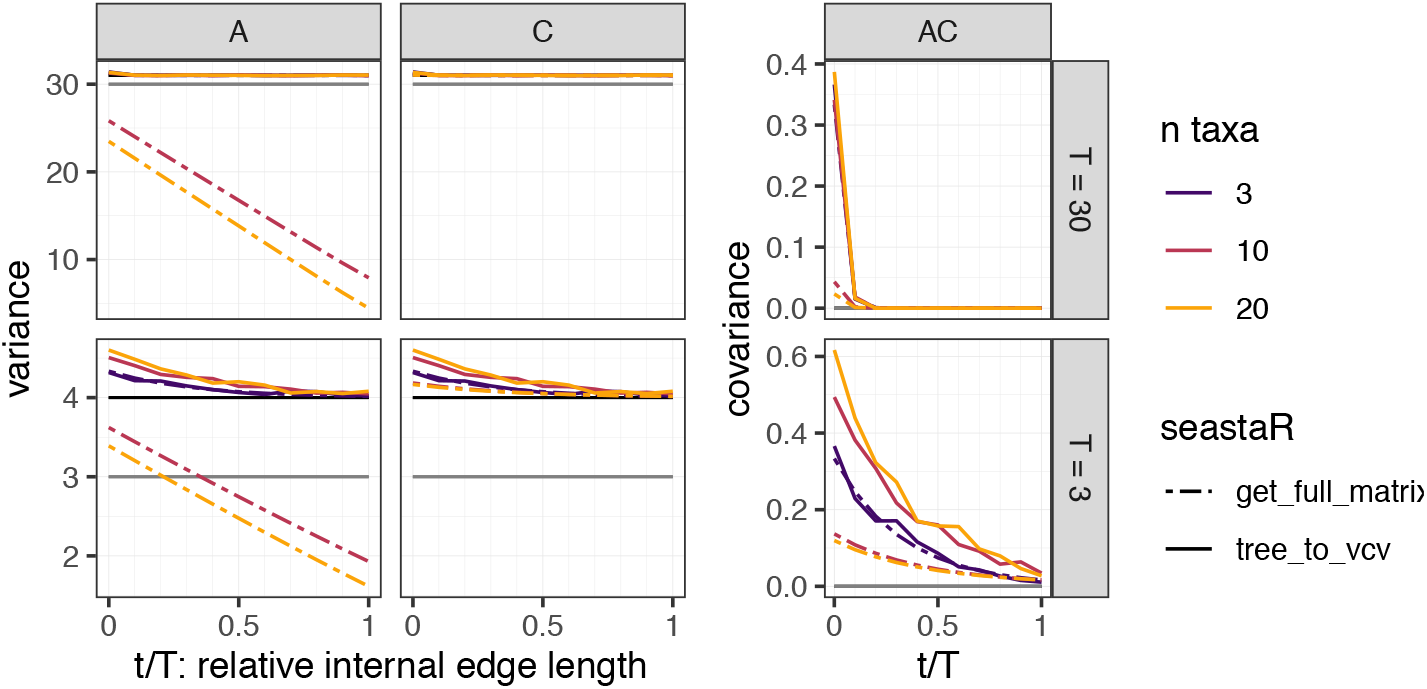
Effect of taxon sampling and clade’s distance to the root on the variance and covariance calculated by various methods on the species tree in Fig. 2 (left) with branch lengths (*T* and *t*) in coalescent units. **Left**: trait variance at *A* and at *C* . **Right**: covariance between the traits at *A* and *C* . Colored dashed lines show *C*^∗^ values from `get_full_matrix`, as in Fig. 3. Colored full lines show values from `trees_to_vcv` on *N*_*g*_ = 1 000 simulated gene trees. The value from the GC model is shown by a black horizontal line (at *T* + 1 = 31 or 4 for the variance, 0 for the covariance). The value from the standard BM model (without ILS) is indicated with a dark gray horizontal line (at *T* = 30 or 3 for the variance, 0 for the covariance).

**Figure S4.**
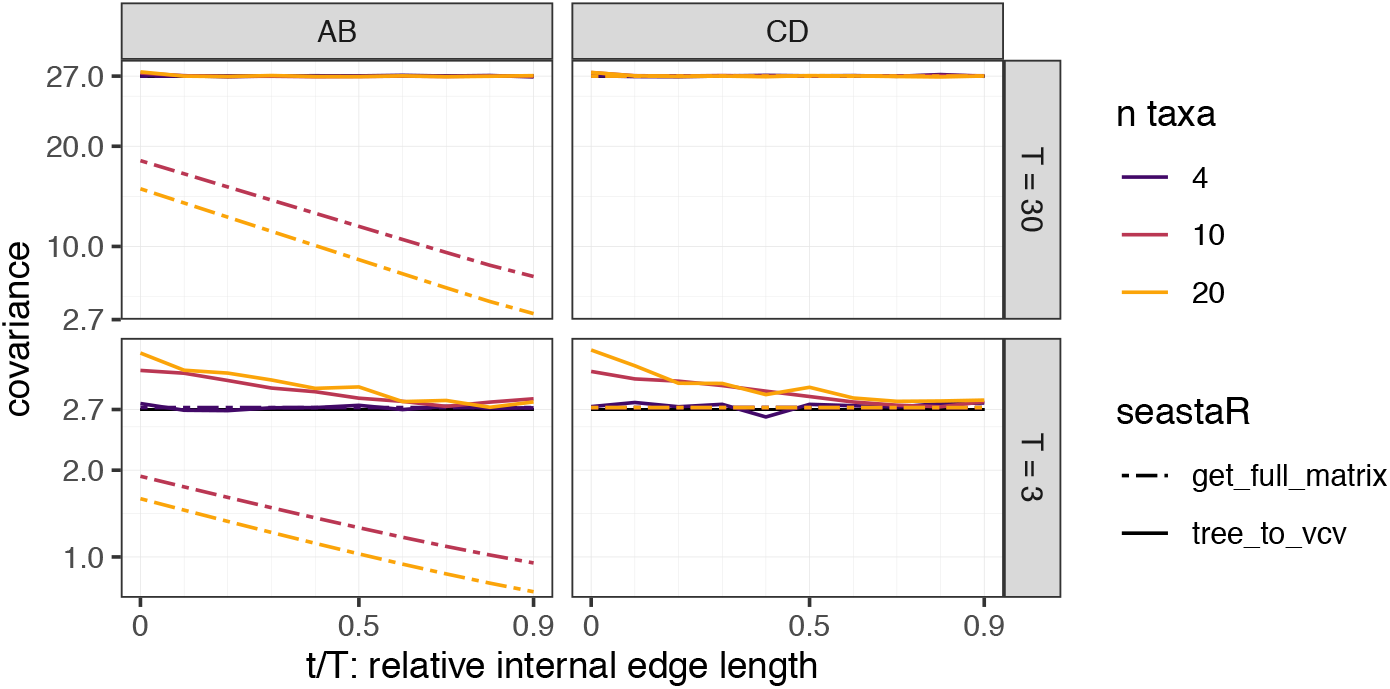
Effect of taxon sampling and clade’s distance to the root on the covariance *C*^∗^ of two sister pairs, on the species tree in Fig. 2, right, with branch lengths (*T* and *t*) in coalescent units, as in Fig. 4. **Left**: trait covariance between *A* and *B*. **Right**: trait covariance between *C* and *D*. The value from the GC model is shown by a black horizontal line. It is identical to the value from the standard BM model (without ILS), at 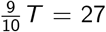 (top) or 2.7 (bottom).

Finally, the two-step procedure using `trees_to_vcv` gives noisy results, especially for larger species trees. This is expected from the procedure itself, which samples a finite number of gene trees (here, *N*_*g*_ = 1 000). Further analyses could simulate more trees, for instance scaling *N*_*g*_ with the number of populations *n*, and `trees_to_vcv` should ideally output standard errors so the user can decide if *N*_*g*_ is sufficiently high, and obtain error bars. Such analyses could however be time consuming, on large phylogenies. Figure S5 shows that the method based on `trees_to_vcv`, and, to a lesser extent, the `get_full_matrix` method, are both much slower than the traditional BM variance calculation method (using `vcv` from the `ape` package) or our GC approach implemented in `phylolm`, and that their timing performances sharply worsen with the number of species. For instance, computing a single covariance matrix for a tree with 20 species takes on average about 1 millisecond for `ape::vcv` and `phylolm` (GC), but about 0.7 second for `get_full_matrix` and around 24 seconds for `trees_to_vcv` (after simulating *N*_*g*_ = 1 000 gene trees).

**Figure S5.**
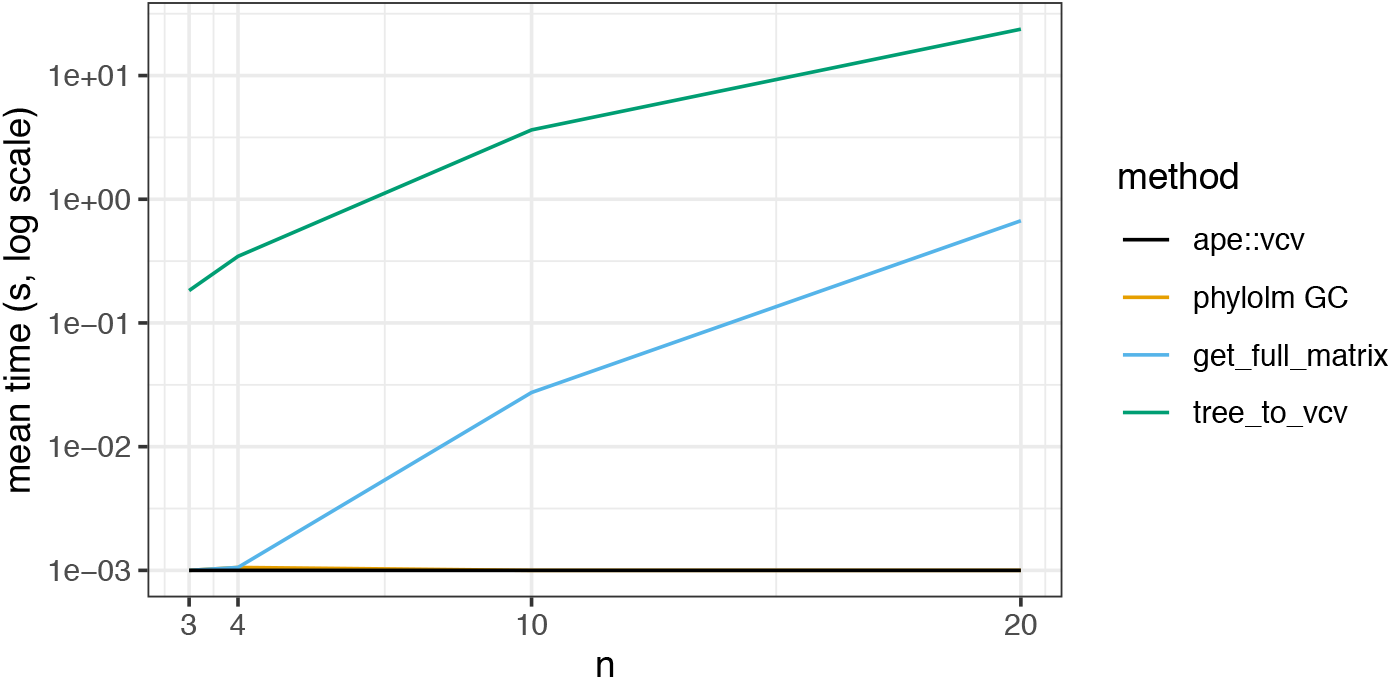
Average time, in seconds, to compute the correlation matrix for one species tree with *n* species. For the method based on `trees_to_vcv`, the time to simulate gene trees is not included. Times below the `R` precision to measure CPU time (1ms) or measured as zeros were arbitrary set to 0.001 seconds.

### D.2. Simulations on a network

The following figures show the estimation accuracy of various parameters of our Gaussian-Coalescent model from simulations. Data were simulated under a 17-taxon network with 3 reticulations, 20 individuals per taxon, and traits controlled by *L* = 100 loci with compound Poisson-Laplace effects (see details in Section 3.2). Branch lengths in the network were rescaled for a total height of *T* = 10 for low ILS, and *T* = 1 for high ILS. The true evolutionary variance rate was fixed at 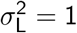. We varied the ancestral trait variance in the root population 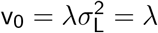.

**Figure S6.**
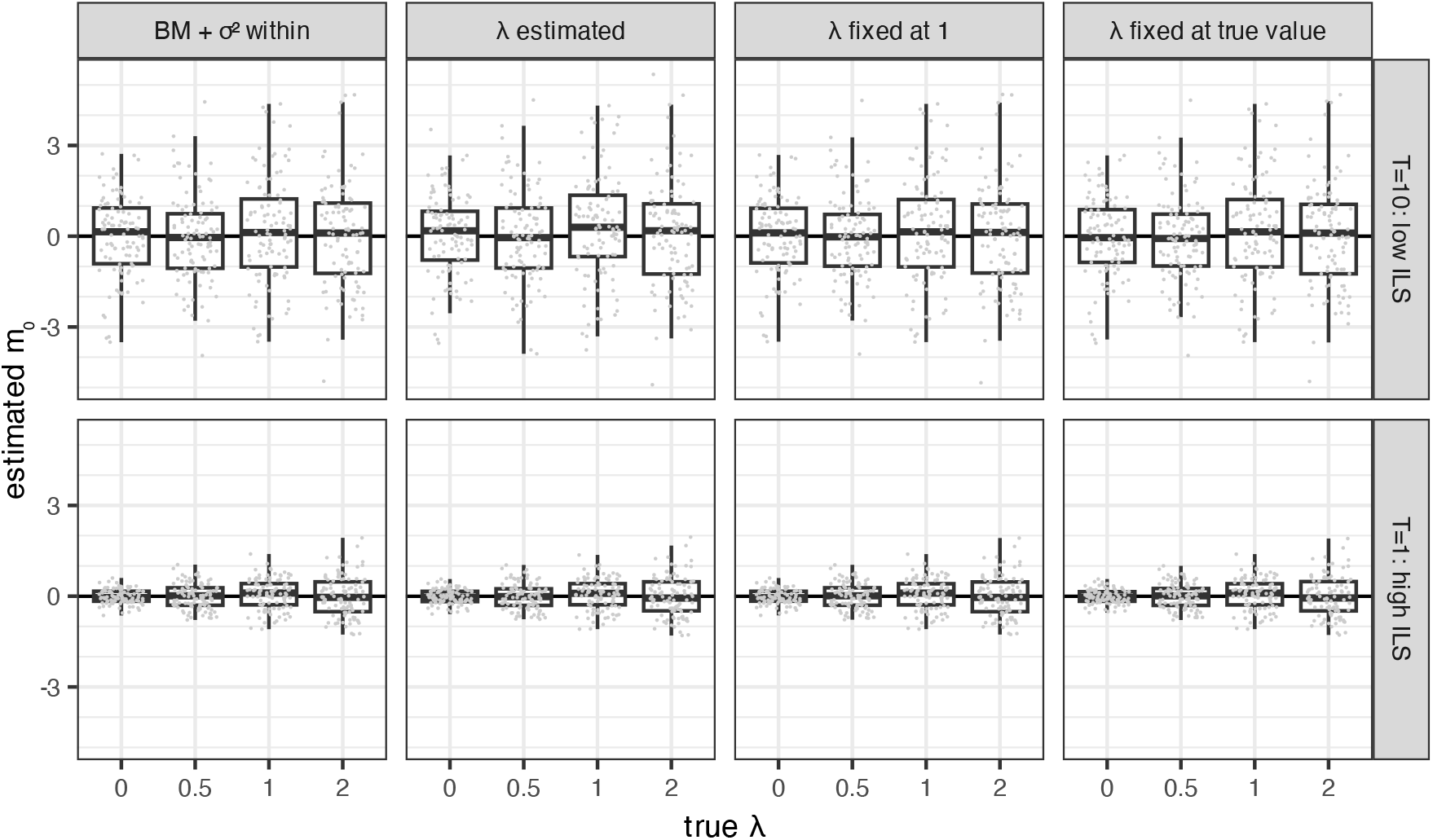
Estimated mean of the ancestral population, whose true value was *m*_0_ = 0. Phylogenetic correlation is higher under low ILS (top) than under high ILS (bottom), resulting in less precision. Under high ILS, estimation error increases with *λ* because it is affected by the root variance v_0_ = *λ*.

**Figure S7.**
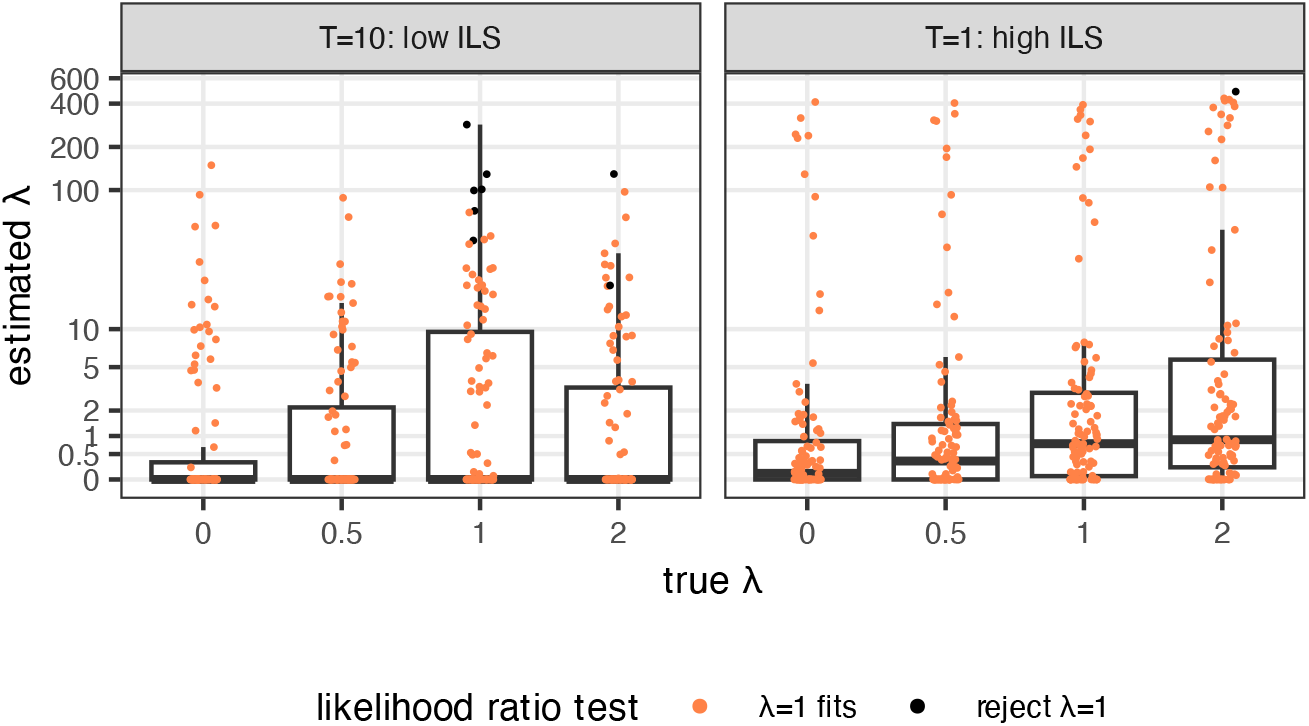
Estimated 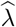, displayed on the log(· + 1) scale, when 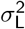 and *λ* were estimated simultaneously. Under low ILS (left), the trait data contain very little information about *λ*. Estimates are slightly more concentrated under high ILS (right). Under both low and high ILS, the null hypothesis that *λ* = 1 is rejected (black dots) very rarely by a likelihood ratio test at the 5% level.

**Figure S8.**
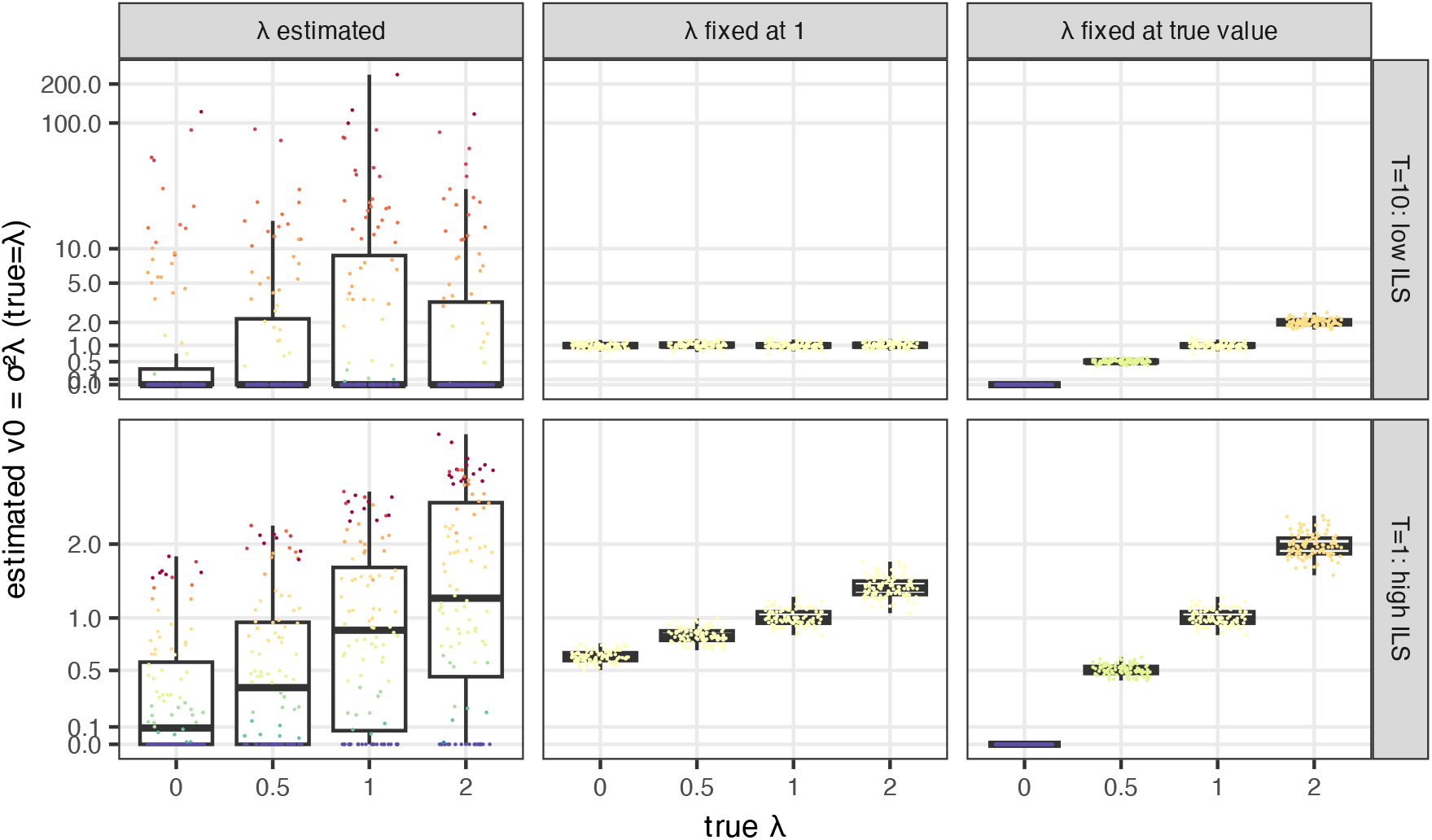
Estimated ancestral state variance 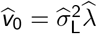, displayed on the log(· + 1) scale. Note that the top row (low ILS) and bottom row (high ILS) use different limits for the vertical axis. If *λ* is known correctly (right), v_0_ is well estimated. But if *λ* is unknown, as is realistic in practice, the trait data contain little information about v_0_, especially under low ILS. Colors indicate the estimated (or fixed) *λ*, with cold (blue) colors for low values and warm (red) colors for large values.

**Figure S9.**
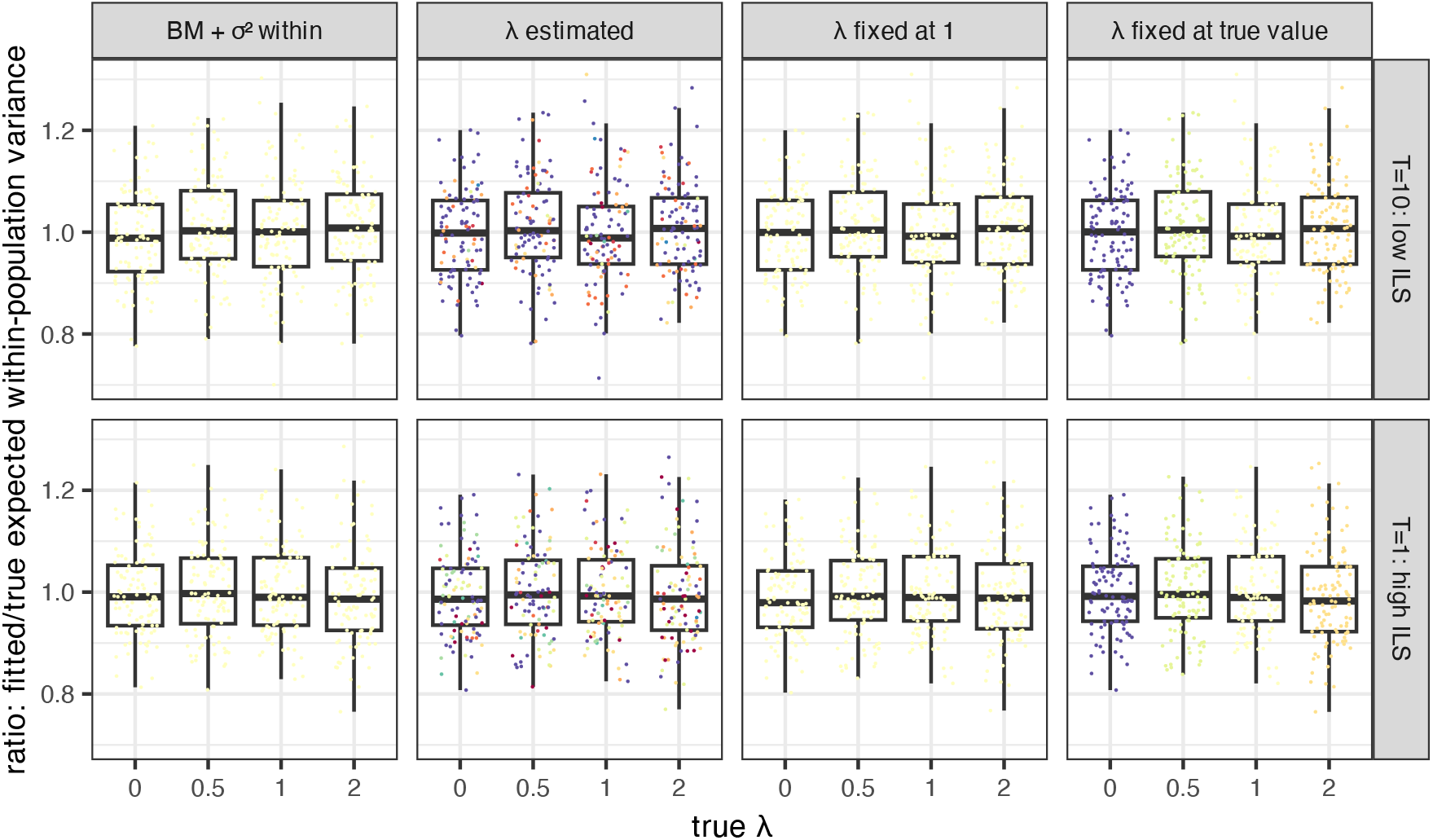
Ratio 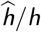, where *h* is the average expected within-species variance: expected variance *H*_*u*_ in population *u*, averaged across all 17 populations. For *u* not below any reticulation, *H*_*u*_ is given in (20). Generally, *H*_*u*_ can be calculated recursively, using the true parameters. When fitting the data with the GC model, 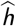 is calculated similarly but using the estimated parameters 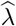 and 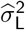. When fitting the data with the BM model, 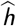 was set to the estimated within-species variance.

### D.3. Simulations on the tomato tree

#### D.3.1. Setting

We reproduced the simulations described in Section 3.2, but using the empirical tomato tree from Section 3.3 instead of the scaled *Polemonium* network. More preciselly, we took the tree that was used in the “full dataset” tomato analysis, with 12 populations and a total height of *T* = 6 coalescent units (Fig. 7), and kept only 3 individual per populations. We then simulated polygenic traits on the tree using the same set-up as described in Section 3.2, except that we simulated *n*_*i*_ = 3 individuals per population.

#### D.3.2. Results

Results on the tomato tree were on par with those obtained under the “low ILS” setting above. The evolutionary variance rate 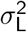 was well estimated by all models (Fig. S10, top row), but with a higher precision for GC models compared to the BM model: the root mean square error (RMSE) for the GC with *λ* estimated, fixed to 1 or fixed to its true value were, respectivelly, equal to 0.127, 0.120 and 0.119, versus 0.269 for the BM. There was again no noticeable gain to knowing the true value of *λ*. The estimate of the mean *m*_0_ in the ancestral root population was unbiased for all methods (Fig. S10, bottom row).

Under the GC model, the estimation of *λ* was again extremely imprecise (Fig. S11). Using a likelihood ratio test at level *α* = 0.05, *λ* = 1 was rejected in only 3 of the 400 replicates in all simulation settings (black dots in Fig. S11). Using AIC, the GC model with fixed *λ* = 1 was favored 80% of the time, followed by the BM model 15% of the time. Estimating *λ* was deemed worthwhile only 4% of the time (Fig. S12).

**Figure S10.**
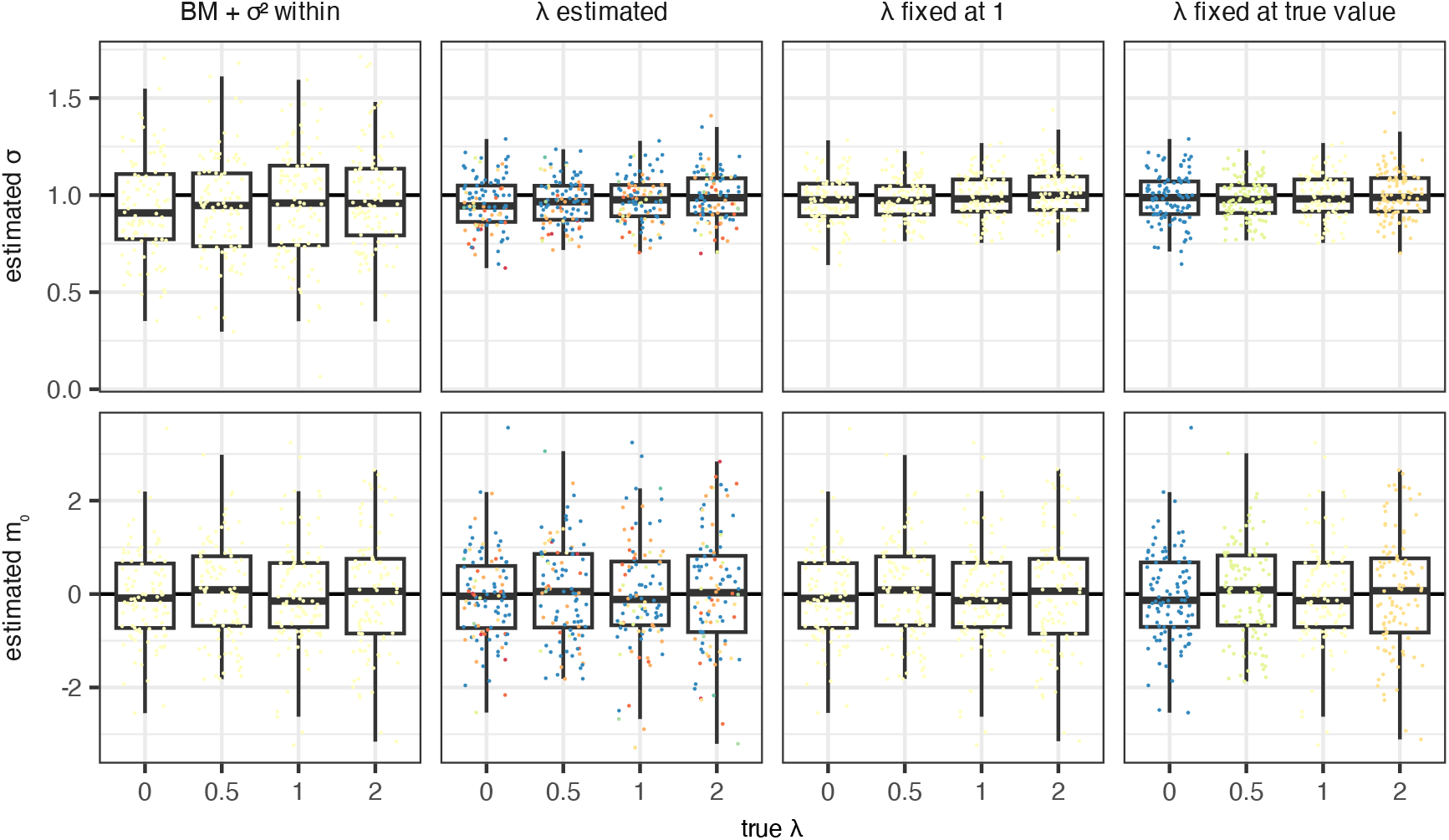
Estimated standard error rate 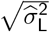 (top) and expectation of the ancestral population 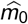 (bottom) from data simulated under the tomato 12-taxon tree, 3 individuals per taxon, and traits controlled by *L* = 100 loci with compound Poisson-Laplace effects. The true parameters were 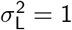 and *m*_0_ = 1. Each column corresponds to one estimation method: the BM model with extra within-population variation, the GC model with *λ* estimated, fixed at 1, or fixed at its true value (from left to right). Colors indicate the estimated or fixed *λ* (or yellow for the BM model), with cold colors for low values and warm colors for large values.

**Figure S11.**
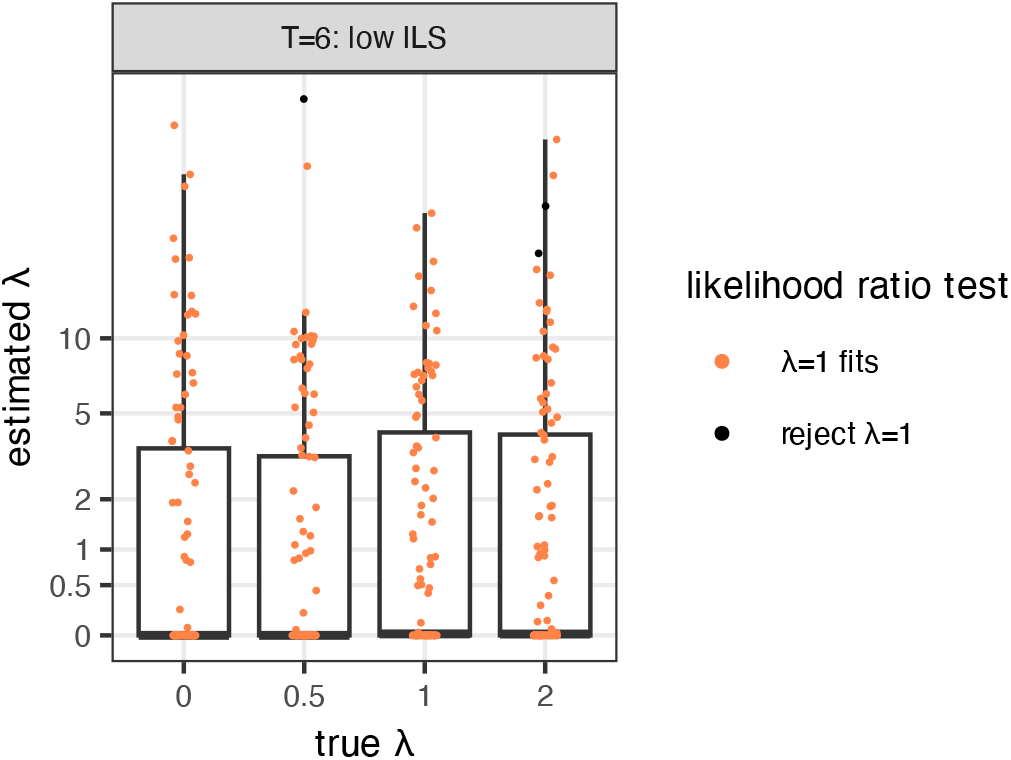
Estimated 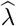, displayed on the log(·+1) scale, when 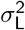 and *λ* were estimated simultaneously. The trait data contain very little information about *λ*. The null hypothesis that *λ* = 1 is rejected (black dots) very rarely by a likelihood ratio test at the 5% level.

**Figure S12.**
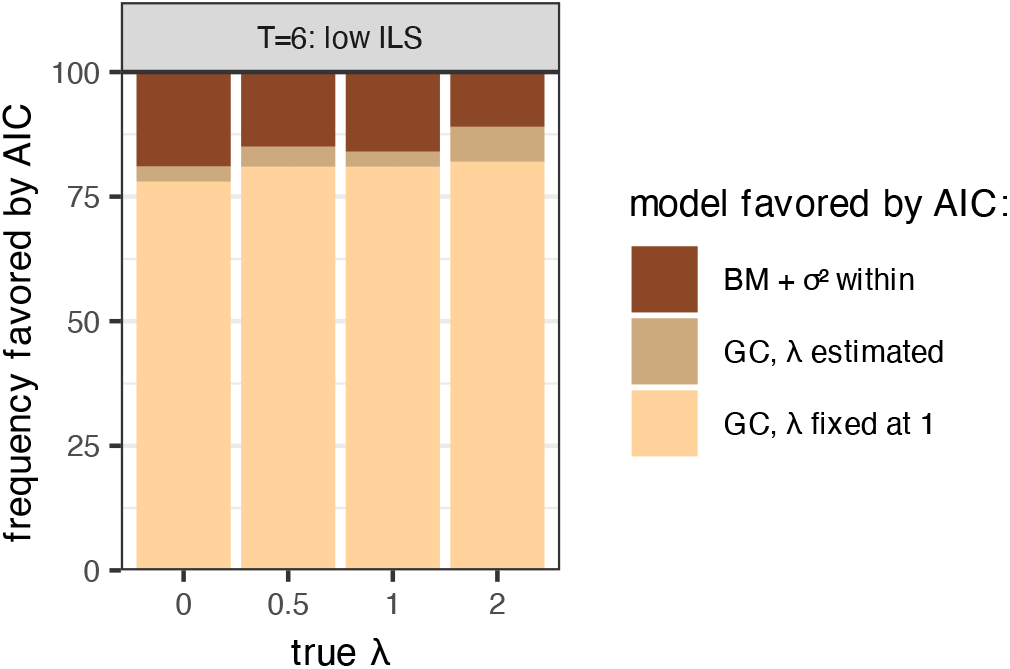
Proportion of times that each model was favored by AIC, out of 100 replicate data sets, when comparing 3 models: Brownian motion with intra-specific variation, GC with *λ* estimated simultaneously with 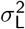, and GC with *λ* = 1 fixed.

## Footnotes

1 See pull request #7 on the `seastaR` repository: https://github.com/larabreithaupt/seastaR/pull/7.

## Notes

https://github.com/cecileane/GCmodel-2026-data-code

https://github.com/JuliaPhylo/PhyloTraits.jl

https://github.com/lamho86/phylolm

https://doi.org/10.5281/zenodo.22083886

### Competing Interest Statement

The authors have declared no competing interest.

### Summary of Updates

Version 3 has been peer-reviewed and recommended by Peer Community In Mathematical & Computational Biology (https://doi.org/10.24072/pci.mcb.100819; Rogers, 2026). This version was updated with the PCI Math. Comp. Biol. badge on the first page, which links to the recommendation.

